# Cytokinin-promoted secondary growth and nutrient storage in the perennial stem zone of *Arabis alpina*

**DOI:** 10.1101/2020.06.01.124362

**Authors:** Anna Sergeeva, Hongjiu Liu, Hans-Jörg Mai, Tabea Mettler-Altmann, Christiane Kiefer, George Coupland, Petra Bauer

## Abstract

Perennial plants maintain their life span through several growth seasons. *Arabis alpina* serves as model *Brassicaceae* species to study perennial traits. *A. alpina* lateral stems have a proximal vegetative zone with a dormant bud zone, and a distal senescing seed-producing inflorescence zone. We addressed the questions of how this zonation is distinguished at the anatomical level, whether it is related to nutrient storage, and which signals affect the zonation. We found that the vegetative zone exhibits secondary growth, which we termed the perennial growth zone (PZ). High-molecular weight carbon compounds accumulate there in cambium and cambium derivatives. Neither vernalization nor flowering were requirements for secondary growth and sequestration of storage compounds. The inflorescence zone with only primary growth, termed annual growth zone (AZ), or roots exhibited different storage characteristics. Following cytokinin application, cambium activity was enhanced and secondary phloem parenchyma was formed in the PZ and also in the AZ. In transcriptome analysis cytokinin-related genes represented enriched gene ontology terms and were expressed at higher level in PZ than AZ. Thus, *A. alpina* uses primarily the vegetative PZ for nutrient storage, coupled to cytokinin-promoted secondary growth. This finding lays a foundation for future studies addressing signals for perennial growth.

**Highlight:** *Arabis alpina* stems have a perennial zone with secondary growth, where cambium and derivatives store high-molecular weight compounds independent of vernalization. Cytokinins are signals for the perennial secondary growth zone.

## Introduction

Perennial plants sustain their life span through several growth seasons and many of these plants are polycarpic and reproduce multiple times (Bergonzi and Albani, 2011). This allows plants to maintain space in their environment to exploit and defend light and soil resources and survive harsh conditions. In contrast, annual plants are usually monocarpic and reproduce only once before they die (Thomas, 2013), a life style of advantage in lowland dry habitats with animal foraging and agriculture in favor of juvenile survival.

Vegetative above-ground shoots of perennial plants may grow in diameter over the years. This production of new tissues in the lateral dimension is termed secondary growth and is initiated by two secondary meristems. The fascicular and interfascicular cambium produces new vasculature in the form of secondary xylem and secondary phloem and corresponding parenchyma, leading to the formation of wood and bast. The cork cambium or phellogen produces outward cork and inward phelloderm to generate new protective outer layers, periderm, with increasing stem diameter. A diverse set of phytohormones and signals influences cambial activity in stems (Fischer et al., 2019). Hormonal profiling across the *Populus trichocarpa* stem identified a specific but interconnected distribution of hormones (Immanen et al., 2016). Cytokinins peaked in the developing phloem tissue, while auxin had its maximum in the dividing cambial cells, and developing xylem tissue was marked by bioactive gibberellin (Immanen et al., 2016). Cytokinins are key regulators of cambium initiation, and cambial activity is affected in cytokinin biosynthetic mutants and by cytokinin supply (Matsumoto-Kitano et al., 2008; Nieminen et al. 2008). Cambial auxin concentration is relevant for activity and maintenance of cambium (Immanen et al., 2016), for example through homeobox transcription factors regulating auxin-dependent cambium proliferation (Suer et al., 2011; Kucukoglu et al., 2017). Gibberellins act in the xylem region promoting xylem cell differentiation and lignification (Denis et al., 2017). Secondary growth is coupled with nutrient allocation. In *Populus*, carbon storage, in the form of triacylglycerols (TAGs), galactose and raffinose, is present in the stem bark and wood tissues (Sauter and van Cleve, 1994; Sauter and Wellenkamp, 1998; Watanabe et al., 2018). Nutrient deposition in storage organs is also affected by plant hormones, and cytokinins may play a dual role in some perennials to direct secondary growth and nutrient storage therein (Hartmann et al., 2011; Eviatar-Ribak et al., 2013).

The perennial life strategy is usually regarded to be ancestral (Hu et al., 2003; Grillo et al., 2009) which is also true for the tribe *Arabideae* within the *Brassicaceae*. Interestingly, in this tribe annuality has evolved multiple times independently from a perennial background in different lineages (Karl and Koch, 2013). This suggests that this complex life style decision may be determined by a limited set of regulatory events. Phylogenetic reconstruction has shown that perennial *Arabis alpina* and annual *Arabis montbretiana* are sister species. Hence, these two taxa have been established as a model system to study the perennial-annual transition (Kiefer et al. 2017; Heidel et al. 2016).

*A. alpina* Pajares (Paj), an accession from Spain, was used to characterize the perennial lifecycle of the species. These plants have above-ground vegetative perennial parts and stems as well as reproductive ones that senesce and die during the growth season (Wang et al. 2009). A characteristic of perennial vegetative stems is the transition from juvenile to adult phase, which results in flowering (Bergonzi and Albani, 2011; Hyun et al. 2017). Competence to flower in perennial *A. alpina* like in annual *Arabidopsis thaliana* can be conferred by the ecologically relevant adaptive trait of vernalization during a cold winter period which suppresses flowering until the action of repressors is released (Kim et al. 2009; Wang et al. 2009). The positions of individual flowers, dormant buds, nonflowering and flowering lateral branches vary along the stems of *A. alpina*. From bottom to top, three types of vegetative (V1, V2, V3) and two inflorescence (I1, I2) sub-zones are distinguished, established during and after vernalization (Lazaro et al. 2018). V1 has full lateral branches resembling the main shoot axis in terms of the zonation pattern. V2 has dormant axillary buds, and the short V3 sub-zone axillary vegetative branches. I1 has lateral branches forming inflorescences, while I2 is an inflorescence and forms individual flowers (Lazaro et al. 2018). The signals and genetic networks triggering the differentiation of V and I zones are not fully understood. *A. alpina PERPETUAL FLOWERING 1 (PEP1*) similar to its orthologue in Arabidopsis, *FLOWERING LOCUS C (FLC*), is a regulator of flowering in response to vernalization and acts as floral repressor (Wang et al. 2009). The *pep1-1* mutant, a derivative of the wild-type accession Paj, shows constant reproduction. Despite lower longevity and the absence of the sub-zonation of the vegetative stem zone, *pep1-1* plants show perennial growth characteristics and have the ability to form continuously new lateral branches until the end of the growth season (Wang et al. 2009; Hughes et al. 2019). Thus, flowering control pathways are regulated by the age control systems to maintain the perennial life style (Hyun et al. 2019). Besides vernalization auxin signals are also involved (Vayssières et al. 2020).

The regulation of nutrient allocation is fundamental for the perennial life cycle to re-initiate growth. In this study, we provide evidence that the proximal vegetative stem zone is characterized by secondary growth where nutrient storage takes place (termed here perennial zone, PZ), irrespective of flowering control by *PEP1*. In contrast, the distal reproductive zone has primary growth (here annual zone, AZ). We furthermore show that cytokinin is effective in initiating secondary growth and we propose a heterochronic model for PZ-AZ transitioning.

## Materials and Methods

### Plant material, growth conditions and plant harvesting

Perennial *Arabis alpina* wild type Pajares (= Paj) accession, perpetual flowering mutant *pep1-1* (Wang et al. 2009) and annual *A. montbretiana* (Kiefer et al. 2017) plants were grown on potting soil (Vermehrungssubstrat, Stender GmbH, Schermbeck, Germany) with a humidity of 70 % in a plant growth chamber (Percival Scientific, CLF Plant Climatics) under controlled long-day (16 h light and 8 h dark, at 20 °C) or short-day (8 h light and 16 h dark, at 4 °C for vernalization) conditions and harvested at stages I, II, III and II’ as described in the Result section and corresponding figures. Plants were fertilized every second week using universal Wuxal (Aglukon, Germany). All experiments were performed with 3-7 biological replicate plants as indicated in the text. For anatomical analysis, 4-11 internode cross sections were prepared per lateral stem sample and stored in 70 % ethanol (Supplemental Figure S1).

Biochemical analysis was conducted with lateral stems from nodes 1 to 5 of the main axis in the case of Paj and *pep1-1* or all lateral stems of *A. montbretiana*. Internodes of one plant were dissected and pooled as one biological replicate, deep-frozen in liquid nitrogen and stored at −80 °C. Entire root systems of single plants were harvested as one root sample and residual soil was removed. Plant samples were freeze-dried, lyophilized and ground to fine powder in a mixer mill (Mixer Mill MM 200, Retsch, Haan, Germany). All biochemical measurements were conducted with the same plant samples (Supplemental Figure S1).

For hormone treatments, *pep1-1* plants were grown as above under long-day conditions for 4 to 5 weeks. The two lateral branches with one elongated internode in a juvenile PZ stage were treated with 300 μl of either 2 mM 6-benzylaminopurine (BAP), 0.5 mM 1-naphthaleneacetic acid (NAA), 4 mM gibberellin GA3 (GA) or 2 mM 1-aminocyclopropane-1-carboxylic acid (ACC) solution (or mock-treated, in 50 % ethanol, mixed with 10 mg lanolin and pasted with a brush along the first internode). The treatment was repeated after 4 days, and internodes were harvested after eight days. AZ treatment was initiated when the plants were 18 weeks old and two elongated internodes of the AZ were formed. Both internodes of five replicate plants were treated with 2 mM BAP solution, as described above. The treatment was performed for 46 days until siliques and uppermost part of the AZ of the control plants started to senesce. Newly formed and already present internodes were treated every fourth day. Harvested AZs were divided into three regions (“top”, “middle” and “bottom”) comprising two to three internodes.

For RNA-sequencing (RNA-seq), 12 *pep1-1* plants were grown per biological replicate. Three biological replicates were used for each examined sampling stage, as described in the text and corresponding figures. Plant samples were immediately deep-frozen in liquid nitrogen, stored at −80 °C and ground to fine powder before use.

### Microscopic analysis

50-100 μm hand-made cross internode sections were incubated for 5-8 minutes in 1 mg/ml fuchsine, 1 mg/ml chrysoidine, 1 mg/ml astra blue, 0.025 ml/ml acetic acid staining (= FCA staining) solution, washed in water and alcohol solutions and mounted. For starch staining, sections were incubated for 5 minutes in 50 % Lugol solution (Sigma-Aldrich GmbH), rinsed in water and mounted. For lipid staining, sections were incubated for 30 minutes in a filtered solution of 0.5 % Sudan IV in 50 % isopropanol, rinsed in water and mounted. Pictures were taken with an Axio Imager 2 microscope (Carl Zeiss Microscopy, Jena, Germany), equipped with an Axiocam 105 color camera (Carl Zeiss Microscopy, Jena, Germany) using bright-field illumination. Images were processed and tissue width quantified via ZEN 2 (blue edition) software (Carl Zeiss Microscopy, Jena, Germany).

### Starch quantification

Starch was quantified enzymatically according to Smith and Zeeman (2006). Briefly, starch was extracted by boiling 10 mg lyophilized plant material in 80 % ethanol. Extracted starch granules were gelatinized at 100 °C. After incubation with α-amyloglucosidase and α-amylase the samples were assayed for glucose in a microplate reader (Tecan Group, Männedorf, Switzerland) using hexokinase and glucose 6-phosphate dehydrogenase to monitor reduction of NAD^+^ at 340 nm (extinction coefficient 6220 l mol^−1^ cm^−1^).

### Triacylglycerol (TAG) quantification

TAGs were obtained by fractionation of glycerolipids and quantified based on their fatty acid contents, normalized to plant sample dry weight (Sergeeva et al.). Briefly, lipids were isolated from dried plant samples according to a modified acidic chloroform/ methanol extraction method for glycerolipids (Hajra et al., 1974; Wewer et al., 2011). TAGs were separated from glycolipid and phospholipid fractions by successive elution with chloroform, acetone/ isopropanol, methanol. Fatty acid methylesters (FAMEs) of the TAG fatty acids or of the total fatty acids from plant samples were analyzed by gas chromatography-mass spectrometry (GC-MS) and resulting peaks integrated for quantification (Sergeeva et al.).

### Protein content determination

10 mg lyophilized plant material was taken up in 200 *μ*l (for stems) or 300 *μ*l (for roots) of Laemmli buffer (2 % SDS, 10 % glycerol, 60 mM Tris-HCl pH 6.8, 0.005 % bromphenol blue, 0.1 M DTT) and heated at 95 °C for 10 minutes. Total protein contents were assayed using the 2-D Quant Kit (GE Healthcare, Little Chalfont, UK), measuring OD_480_ in a microplate reader (Tecan Group, Männedorf, Switzerland) using a bovine serum albumin mass standard curve.

### Determination of carbon (C) and nitrogen (N)

For C and N quantification, 2 mg of lyophilized plant material, packed into tin containers, was applied to elemental analysis isotope ratio mass spectrometry (EA-IRMS) (Elementar Analysensysteme, Germany). C/N ratios were calculated from C and N values.

### RNA isolation

Total RNA from plant samples was isolated using the RNeasy Plant Mini Kit (Qiagen, Hilden, Germany), including DNase I digestion via the RNase-Free DNase Set to remove traces of genomic DNA (Qiagen, Hilden, Germany). Quality and quantity of the isolated RNA was examined with the Fragment Analyzer (Advanced Analytical Technologies GmbH, Heidelberg, Germany). All samples had a suitable RNA quality number above 7.0 (RQN mean = 8.9).

### RNA sequencing and analysis

RNA-seq analysis was conducted for three biological replicates per sample. For gene expression profiling, libraries were prepared using the Illumina TruSeq^®^ Stranded mRNA Library Prep kit. Prepared libraries were sequenced on the HiSeq3000 system (Illumina Inc. San Diego, CA, USA) with a read setup of 1 x 150 bp and an expected number of 28 Mio reads.

Using the latest publicly available versions of the *A. alpina* genome assembly (Arabis_alpina.MPIPZ.V5.chr.all.fasta and Arabis_alpina.MPIPZ.V5.chr.all.liftOverV4.v3.gff3, both downloaded from arabis-alpina.org), 34.220 coding sequences (CDS) were assembled. Closest *A. thaliana* orthologs were determined by blasting *A. alpina* CDS assemblies against the most recent version of the *A. thaliana* proteome sequences (TAIR10_pep_20101214_updated; downloaded from TAIR) using the *blastx* algorithm of the Blast+ suite (Altschul et al., 1990) with an E-value threshold of < 1E5. Among multiple resulting hits, the Arabidopsis proteins with the highest bit score followed by the lowest E value were accepted as the closest orthologs if the percentage of identity was > 25 % of the aligned amino acids (Doolittle, 1986) and the E-value was < 1E-15. Single end reads were trimmed with *trimmomatic* (Bolger et al., 2014) and the quality of the trimmed reads was evaluated with *fastqc* (Andrews, 2010). Using *kallisto* (Bray et al., 2016), the trimmed reads were mapped to the *A. alpina* CDS assemblies and quantified, which resulted in estimated counts and transcripts per million (TPM) values per gene. As a control, the trimmed reads were mapped to the genome assembly with *hisat2* (Kim et al., 2015) and transcripts were quantified with *htseq-count* (Anders et al., 2015) using a gene transfer format (*.gtf) file that was generated from the general feature format (*.gff3) file. The resulting counts were transformed to TPM and used for Principal Component Analysis (PCA, R: *prcomp*) and hierarchical clustering (HC, R: *dist, hclust*). Estimated counts were used for statistical analysis using *edgeR* (Robinson et al., 2010; McCarthy et al., 2012). Resulting p-values were adjusted with the Bonferroni method. Fold-change values of gene expression were calculated in pairwise comparisons between samples and pooled PZ and AZ samples. The above-described calculations were performed with the values obtained from *kallisto* and with those obtained from *hisat2* and *htseq-count*. Genes were accepted as differentially regulated if p < 0.01 with both mapping and quantification methods. Graphs were produced with TPM obtained by *kallisto*.

Gene ontology (GO) analysis was carried out using *topGO* (Alexa and Rahnenfuhrer, 2010) with the closest *A. thaliana* ortholog Locus ID’s (AGI code) using the latest publicly available *A. thaliana* GO annotations (go_ensembl_arabidopsis_thaliana.gaf; downloaded from Gramene) applying Fisher’s exact test. GO terms were enriched with p < 0.05.

### Gene expression by reverse transcription-qPCR

Total RNA was reverse-transcribed into cDNA using oligo dT primer and diluted cDNA used for qPCR, as described (Ben Abdallah and Bauer, 2016). Absolute normalized gene expression values were calculated based on mass standard curve analysis and reference gene normalization (Stephan et al. 2019). Primers for qPCR are listed in Supplemental Table S1.

### Statistical analysis

R software was used to perform statistical analyses by one-way analysis of variance (ANOVA) in conjunction with Tukey’s HSD (honest significant difference) test (α = 0.05). Significant differences with p < 0.05 are indicated by different letters.

## Results

### Lateral stems of *A. alpina* plants have a perennial secondary growth and annual primary growth zone

To investigate nutrient storage in *A. alpina* stems, we first inspected the anatomy of stem internode cross sections. Since nutrient storage might be coupled with signals for vernalization, we included besides vernalization-dependent wild-type Pajares (Paj) also its mutant derivative *perpetual flowering 1-1* (*pep1-1*), that flowers independently of vernalization. We performed comparisons of perennial lines to annual sister species *A. montbretiana* which does not require vernalization, either. We focused on internodes of the lateral branches, formed in the V1 subzone of the main axis at the lower first to fifth nodes. Sub-zonation into three V zones (V1, V2 and V3) is present in Paj, but absent in *pep1-1*. Instead, *pep1-1* has a single V zone, V1, and all axillary buds develop into flowering branches (Wang et al., 2009; Vayssières et al. 2020). Thus, by investigating lateral branches from V1 we were able to study the role of *PEP1* in stem development and nutrient allocation.

The anatomy of these lateral branches changed from the vegetative (V) to inflorescence (I) zone. Paj V internodes displayed secondary growth with cambium and cork cambium activities. Since advanced secondary growth is a perennial trait, we termed this zone “perennial zone” (“PZ”, Figure 1). I zone internodes had only primary growth characteristics in Paj, and accordingly, we termed this zone “annual zone” (“AZ”, Figure 1).*pep1-1* showed a PZ and an AZ as Paj, while *A. montbretiana* lateral stems displayed the AZ pattern (Figure 1).

**Figure 1:**
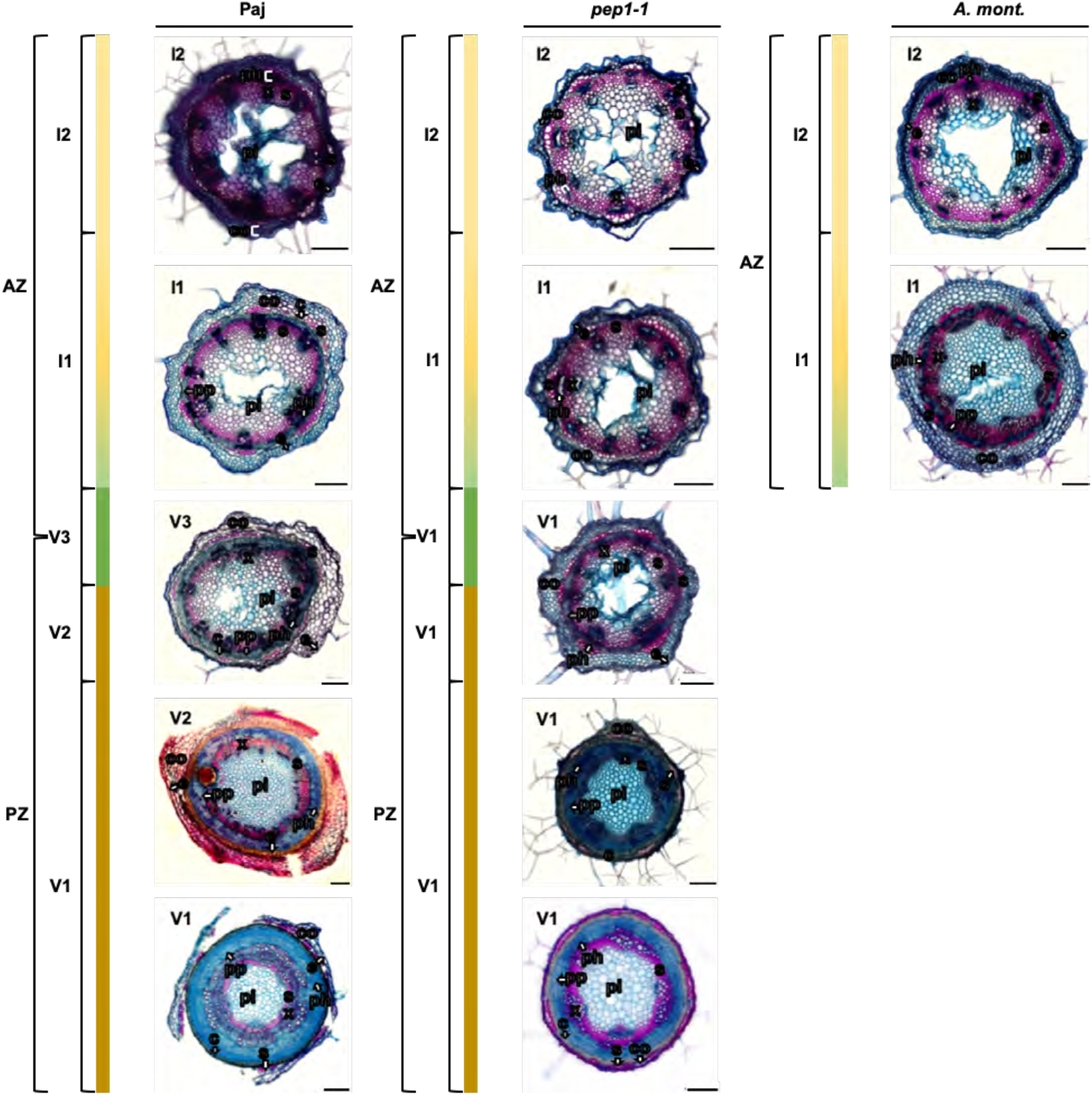
Overview of lateral stem internode anatomy of perennial and annual *Arabis* plants. 100 μm cross sections of lateral stems formed at the lower nodes of main stems from perennial *A. alpina* Pajares (Paj, wild type), its *perpetual flowering1-1* mutant derivative (*pep1-1*) and annual *A. montbretiana (A. mont.*). Plants were harvested at different growth stages (see Figure 2, red triangles indicate positions of cross sections). A „perennial zone” (PZ) with secondary growth characteristics is found at the proximal side of lateral branches, corresponding to the vegetative zone (V1, V2, described for main stems in Lazaro et al. 2018), while an „annual zone“ (AZ) with primary growth overlaps with inflorescence zone (I1, I2, described in main stems in Lazaro et al. 2018). PZ and AZ are separated by a transition zone (V3, described for main stems in Lazaro et al. 2018). Anatomy, investigated following FCA staining, in blue, staining of non-lignified cell walls (parenchyma, phloem, meristematic cells), in red, staining of lignified cell walls and greenish, staining of suberized cell walls (xylem, sclerenchyma, cork). Abbreviations used in microscopic images: c, cork; co, cortex; e, epidermis; ph, phloem including primary phloem, secondary phloem, phloem parenchyma; pi, pith; pp, secondary phloem parenchyma; s, sclerenchyma; x, xylem including primary xylem, secondary xylem, xylem parenchyma. Arrows point to respective tissues. Scale bars, 200 μm.

Next, we inspected lateral stem anatomy more closely at specific growth stages during development of the PZ and AZ, namely at stage I (after 8 weeks, before vernalization), at stages II and III (after 20 and 35 weeks including vernalization) and at stage II’ (after 20 weeks without vernalization) (Figure 2, see Supplemental Figure S2 for detailed descriptions of plant growth). At stages I and II, lateral branches had vegetative character (Figure 2B) and all elongated internodes corresponded to PZ with beginning secondary growth, similar in Paj and *pep1-1* (PZ in Figure 3A, Figure 3B). At stage I (Figure 3A), epidermis and cortex were separated from the central pith by a ring of vascular bundles. Cambium was narrow, surrounded by few layers of secondary phloem. We noted a ring of cork cambium below the sclerenchyma cap in vascular bundles and below the outer cortex in the interfascicular region. Cork cells were formed already in some regions (note that this tissue became more evident at later stages). At stage II (Figure 3B), cork formation underneath the outer cortex had progressed to a multi-cell-layer-thick ring in the PZ. Cortical and epidermal cells were compressed and showed lignification (appeared reddish by FCA staining, Figure 3B). At stage III, lateral branches of Paj flowered and formed green siliques, and those of *pep1-1* were in a more advanced reproductive state and already senescent (Figure 2B). Secondary growth was further progressed in the PZ (PZ in Figure 3C). The lower lateral stem internodes of Paj and *pep1-1* had hardly any outer cortex left, and if present, the cortex cells were compressed. Frequently, sclerenchyma, outer cortex and epidermis peeled off as fibers during preparation. The cambium had formed a large ring of secondary phloem and secondary xylem with parenchyma. The upper lateral stem internodes of the AZ showed the annual growth pattern (AZ in Figure 3C). Interestingly, in the AZ, interfascicular cambium and a ring of phloem parenchyma adjacent to the fascicular phloem were found in both Paj and *pep1-1* (for Paj visible in stage III, Figure 3C, but for *pep1-1* this was only visible between stages II and III as shown, Supplemental Figure S3A). This interfascicular cambium and phloem parenchyma were no longer visible at later stages of AZ development, but instead sclerenchyma tissue had formed, suggesting that cambium and phloem parenchyma had transformed into sclerenchyma in AZ (for *pep1-1* visible at stage III, Figure 3C, but for Paj visible only later than stage III as shown in Supplemental Figure S3B). The pith regions became hollow in the two lines, and upper stem regions began to senesce. The border of PZ and AZ coincided with a transition region with short internodes. Phloem parenchyma and cork gradually became narrower, until they were no longer apparent (Supplemental Figure S3C, S3D). At stage II’, lateral branches of Paj retained vegetative character and consisted only of PZ internodes (Figure 2B). Contrary to that, lateral branches of *pep1-1* formed the AZ and flowered. Secondary growth was advanced in the PZ, comparable to stage III (PZ, Figure 3D). The upper AZ (only present in *pep1-1* but not in Paj) showed again primary growth (AZ, Figure 3D). *A. montbretiana* plants formed lateral branches and were marked by green siliques (Figure 2B). Lateral stems of *A. montbretiana*, only available at stage II’, showed primary growth (AZ in Figure 3D). Only in a very thin zone close to the main stem secondary growth was apparent in the lateral stems of *A. montbretiana*, presumably necessary for stability (Supplemental Figure S3E).

**Figure 2:**
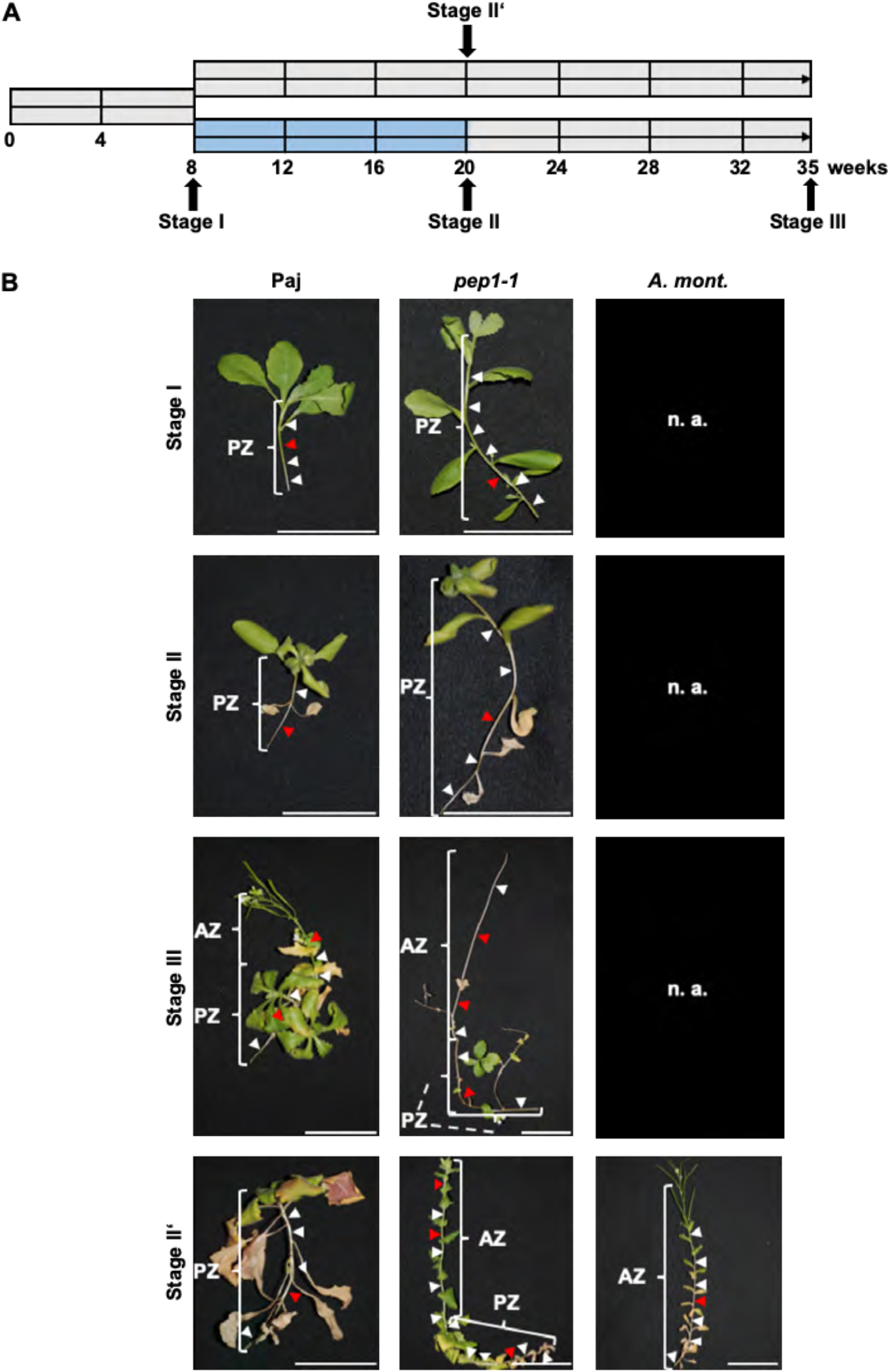
Plant growth and morphology (A) Plant growth scheme and harvesting stages. Time scale of plant growth under, grey color, long-day conditions at 20°C, blue color, short-day conditions at 4°C (vernalization); harvesting stages I, II, III or alternatively II’. (B) Lateral stem morphology at different growth stages. Representative photos of lateral stems from lower nodes of the main axis at stage I, II, III and II’; perennial *A. alpina* Pajares (Paj, wild type), its *perpetual flowering1-1* mutant derivative (*pep1-1*) and annual *A. montbretiana (A. mont.*); for whole plant morphology (Supplemental Figure S2). Red triangles, positions of cross sections in Figure 3, similar as shown in Figure 1; white triangles, other regions examined by microscopy; n. a., lateral stems not present at this stage. Scale bars, 5 cm.

**Figure 3:**
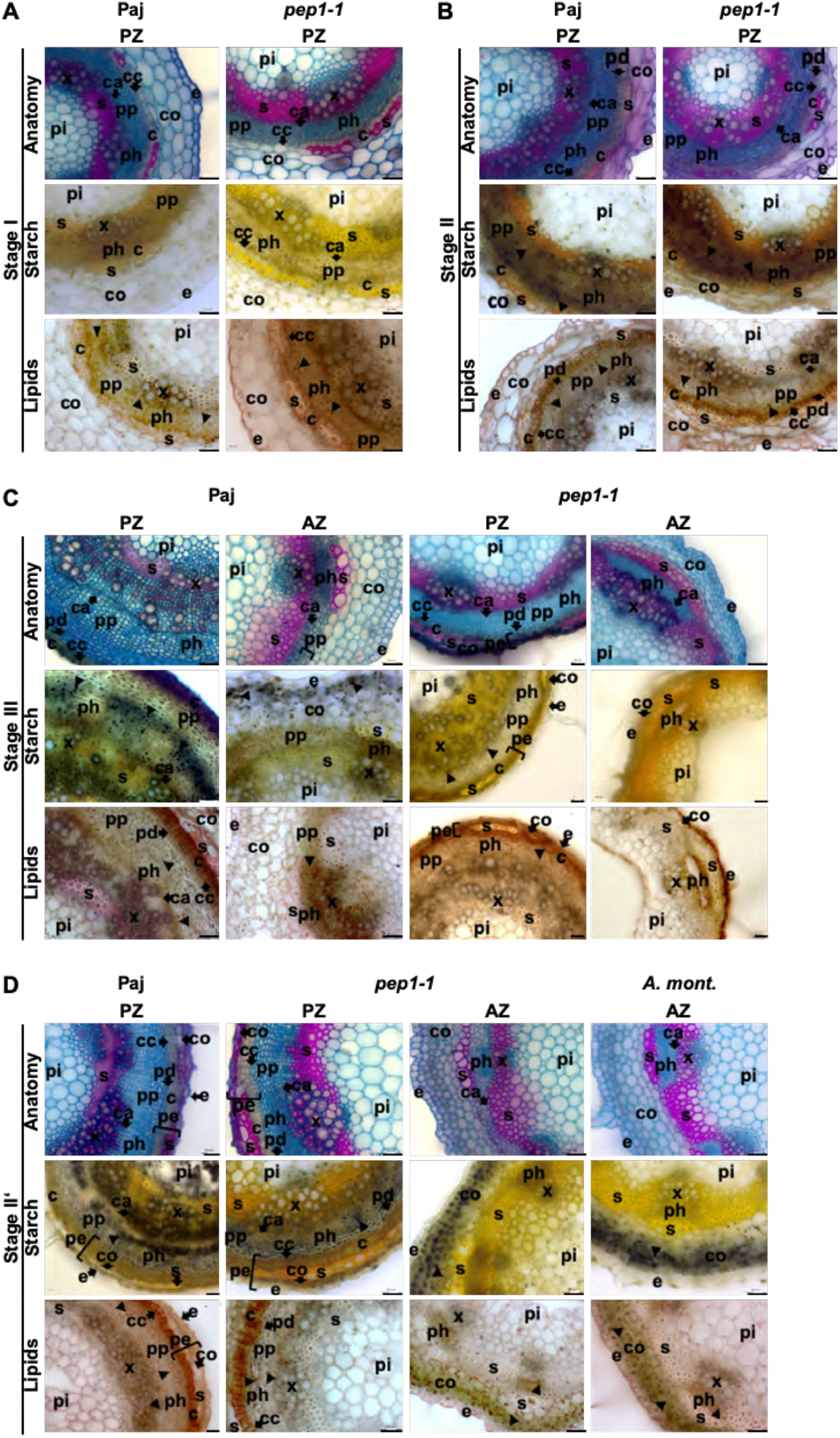
Anatomy, starch and lipid body staining of lateral stem internodes. Representative lateral stem internode cross sections at (A), stage I; (B), stage II; (C), stage III; (D), stage II’, prepared from perennial *A. alpina* Pajares (Paj, wild type), its *perpetual flowering1-1* mutant derivative (*pep1-1*) and annual *A. montbretiana* (*A. mont.*) in the perennial zone (PZ) and annual zone (AZ). Anatomy, investigated following FCA staining, in blue, staining of non-lignified cell walls (parenchyma, phloem, meristematic cells), in red, staining of lignified cell walls and greenish, staining of suberized cell walls (xylem, sclerenchyma, cork). Starch, visualized following Lugol’s iodine staining, dark violet-black color. Lipids, detected by Sudan IV staining, orange pinkish color of lipid bodies and yellowish color of suberized and lignified cell wall structures. Black triangles, starch and lipid bodies. Abbreviations used in microscopic images: c, cork; ca, cambium; co, cortex; e, epidermis; pd, phelloderm; pe, periderm; ph, phloem including primary phloem, secondary phloem, phloem parenchyma; pi, pith; pp, secondary phloem parenchyma; s, sclerenchyma; x, xylem including primary xylem, secondary xylem, xylem parenchyma. Arrows point to respective tissues. Scale bars, 50 μm.

Taken together, the characteristic zonation into an upper reproductive and senescent annual growth zone, the AZ, and a lower vegetative perennial growth zone, the PZ, with secondary growth is a property of perennial *Arabis alpina* Paj and *pep1-1*, not affected by perpetual flowering and vernalization. This zonation was absent in the annual *A. montbretiana*.

### Starch and lipids are stored in secondary growth tissues of the PZ

Secondary growth in the PZ of lateral stems might be linked to nutrient storage. We investigated the deposition of C storage biopolymers starch and lipids (triacylglycerols, TAGs), by staining of tissue sections and biochemical quantification (Figure 3, Figure 4).

**Figure 4:**
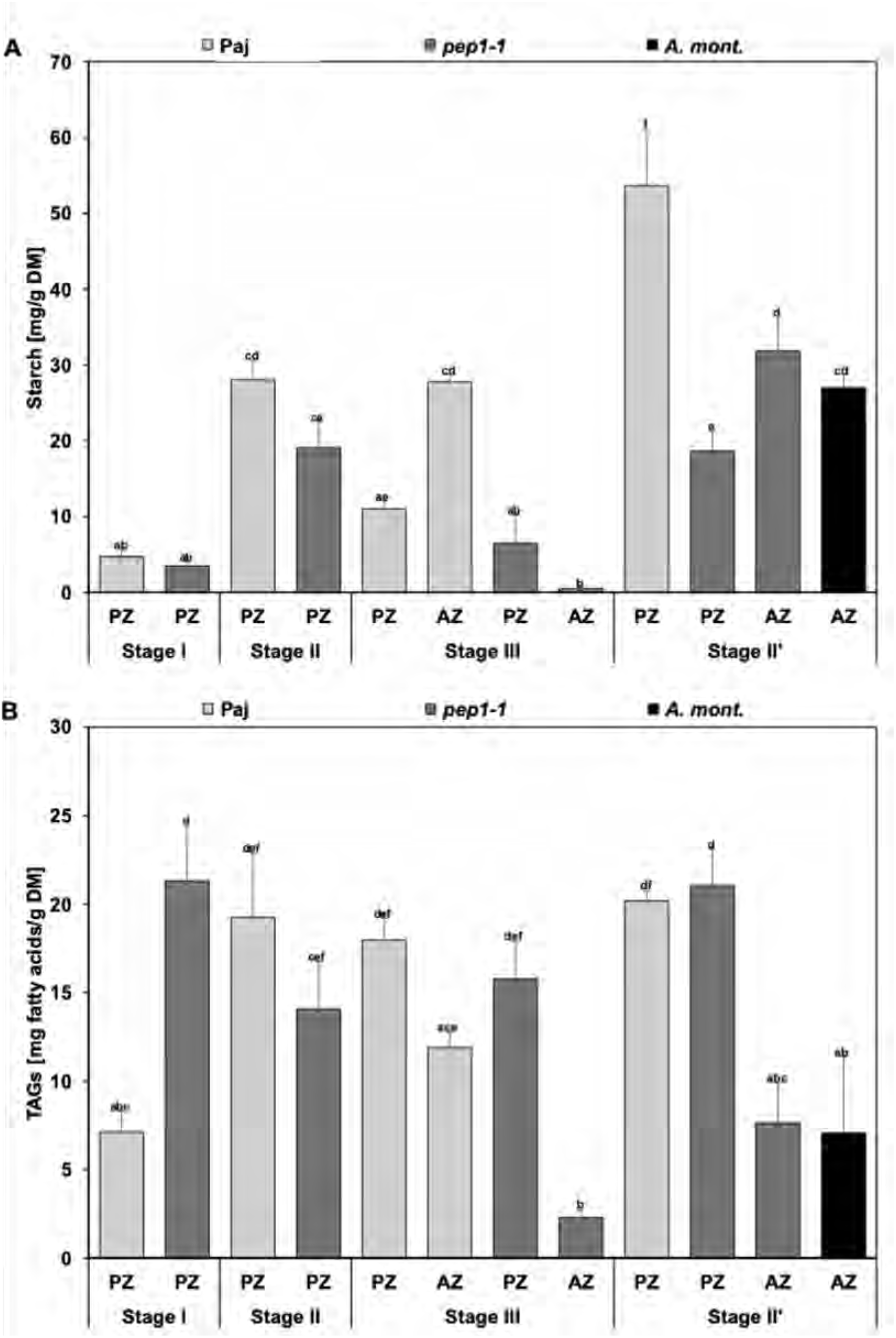
Contents of C-containing storage compounds in lateral stem internodes. Stem diagrams represent contents per dry matter (DM) of (A), starch, and (B), triacylglycerols (TAGs). Lateral stem internodes were harvested at stages I, II, III and II’, corresponding to perennial zone (PZ) and annual zone (AZ) of *A. alpina* Pajares (Paj, wild type), its *perpetual flowering1-1* mutant derivative (*pep1-1*) and annual *A. montbretiana* (*A. mont.*). Data are represented as mean +/- SD (n = 3-7). Different letters indicate statistically significant differences, determined by one-way ANOVA-Tukey’s HSD test (p < 0.05).

Starch was hardly detectable at stage I in PZ in Paj and *pep1-1* (Figure 3A), confirmed by starch contents below 5 mg/g dry matter (Figure 4A). At stage II, starch was formed in the ring of secondary phloem parenchyma in Paj and *pep1-1* (Figure 3B). Starch contents were between 20 and 30 mg/g dry matter (Figure 4A). At stage III, starch was also present in secondary phloem parenchyma of the PZ in Paj and *pep1-1* and contents ranged between 7-12 mg/g dry weight (Figure 3C, 4A). In the AZ, starch was detected in the outer cortex of Paj, but hardly in *pep1-1* (Figure 3C). Indeed, starch contents were higher in the AZ of Paj compared to *pep1-1* (28 mg/g in Paj compared to 0.6 mg/g dry weight in *pep1-1*, Figure 4A). Since AZ development was advanced in *pep1-1* compared to Paj, we figured that starch was consumed and degraded with progression of senescence in the AZ. In stage II’, starch was found deposited in the secondary phloem parenchyma ring of the PZ in Paj and *pep1-1*, while it was confined to the outer cortex of the AZ in *pep1-1* and *A. montbretiana* (Figure 3D). Starch contents were high in stage II’ samples ranging from 18 to 32 mg/g dry matter in *pep1-1* and *A. montbretiana*, and up to even 54 mg/g dry matter in the PZ of Paj (Figure 4A). Significant differences with regard to starch contents between PZs of Paj and *pep1-1* at stage II’ and between PZs of Paj at stage II’ and III not only support the storage property of PZ, but also indicate the active turn-over of stored starch ressources during flowering.

Lipid staining patterns were similar but not identical with starch staining. The main difference was that in the PZ, lipid bodies were found in all stages in the secondary phloem parenchyma, and in addition also in the secondary phloem, secondary xylem and in the cambium (Figure 3A-D). In the AZ, lipid bodies were present in the outer cortex and in addition in the phloem and cambium of vascular bundles in Paj, *pep1-1* and *A. montbretiana* (Figure 3C, 3D). TAG contents did not vary significantly between the PZ of different stages and lines, except at stage I in Paj, where it was lowest in accordance with the presence of lipid bodies in the developing secondary parenchyma (Figure 4B). In all AZ samples, TAG contents were lower than in the corresponding PZ samples, in agreement with anatomical observations (Figure 4B).

In summary, the tissues derived from secondary growth play a predominant role in C storage in the form of starch and TAGs in PZ, irrespective of the flowering status and vernalization.

### Carbon storage is reflected by carbon/nitrogen (C/N) ratio

An elevated C and C/N ratio is an indicator for carbon storage. Total C contents of all stem samples ranged between 360 and 440 mg/g dry matter (Figure 5A). In the PZ, significant increases of the C contents were noted from stage I to stages II and II’ and then remained rather constant. At stage III a lower C content was found in the AZ compared to the PZ, which was significant in *pep1-1* (Figure 5A). Total N contents developed in an opposite pattern to the C contents and decreased significantly from 20-25 mg/g dry mass at stage I to 5-10 mg/g dry mass at stages II and II’ and then remained at a constant low level in stage III (Figure 5B). This finding speaks against usage of N storage compounds. Total protein contents were highest with 110-140 mg/g dry mass in stages I and II and in PZ of Paj at stage III while they were lower and varied between AZ and PZ and between lines without apparent pattern at stages III and II’ (Figure 5C). Perhaps the lower N contents indicate a lower metabolic activity of the fully developed PZ. The C/N ratios were lowest at stage I and increased in stages II, III and II’, however, a distinction of PZ and AZ was not possible and noticeable differences between the lines were not observed (Figure 5D). The C storage capacity of the PZ and possible turn-over of C storage compounds during flowering is reflected by C/N ratio of the PZs of Paj at stage III and II’. The C/N ratio of the PZ retaining vegetative character at stage II’ was significantly higher than that of the PZ corresponding to flowering at stage III. As a comparison to stems, roots may also serve as a C storage reservoir, however, in this case TAGs would be primary C storage products but not starch (Supplemental Figure S4).

**Figure 5:**
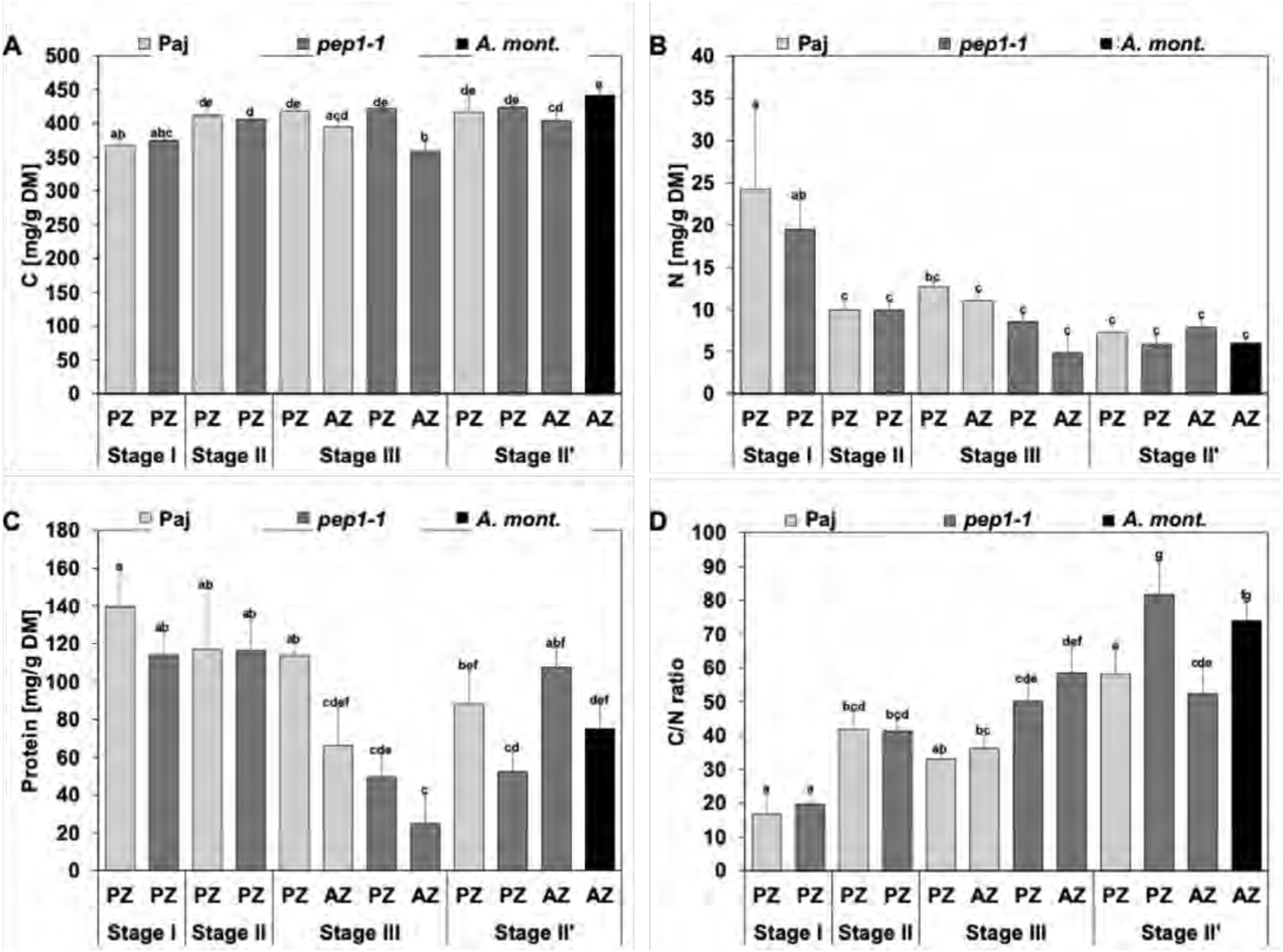
Carbon, nitrogen and protein contents in lateral stem internodes. Stem diagrams represent contents per dry matter (DM) of (A), carbon (C), (B), nitrogen (N), (C), protein, (D), C/N ratios. Lateral stem internodes were harvested at stages I, II, III and II’, corresponding to perennial zone (PZ) and annual zone (AZ) of *A. alpina* Pajares (Paj, wild type), its *perpetual flowering1-1* mutant derivative (*pep1-1*) and annual *A. montbretiana* (*A. mont.*). Data are represented as mean +/- SD (n = 3-7). Different letters indicate statistically significant differences, determined by one-way ANOVA-Tukey’s HSD test (p < 0.05).

Taken together,, the total C content increased slightly with the progression of the development of the PZ. This slight increase reflects storage of starch and TAGs but also may comprise the increases in cell walls.

### Hormones affect secondary growth in lateral stems

We tested effects of hormone application to internodes in a juvenile PZ stage prior to visible secondary growth. First, we applied synthetic 6-benzylaminopurine (BAP) for eight or 46 days to PZ and AZ internodes respectively. In the PZ, cambium and phloem parenchyma regions were increased in width (Figure 6A, 6B, “Anatomy”). The increase in width corresponded to a doubling of cambium cells from two to about four cells and additional one or two cells of phloem parenchyma in the fascicular and interfascicular regions (Figure 6A-D). Interestingly, the increase in width was also found in the newly developed second internode that had not been treated with BAP directly, showing that a cytokinin-induced signal acted in the adjacent internode (Figure 6C, 6D). Similar to the control plants, starch did not accumulate in the secondary phloem parenchyma of the treated plants. Patterns of lipid bodies were similar in treated and control stems (Figure 6A, 6B, “Starch”, “Lipids”). After 46 days of BAP treatment, the AZ internodes also increased in width (Supplemental Figure S5B). Additionally, flowering was delayed and a second zone with short internodes, as found in the PZ-AZ transition zone, was formed two to three internodes above the first (Supplemental Figure S5B). Cambium and phloem parenchyma regions were considerably increased in width in the AZ (Figure 7A, 7B, “Anatomy”, 7C, 7D). Secondary phloem parenchyma was formed in the interfascicular regions of “Top” and “Middle” areas of the AZ. In the control, the corresponding areas would differentiate into sclerenchyma (Figure 7B, “Anatomy”: “Top” and “Middle”, 7D). Furthermore, cork cambium and cork were formed in the “Middle” area in the fascicular and interfascicular regions of the treated stems (Figure 7A, 7B). A difference with regard to starch accumulation was observed only for the “Bottom” area (Figure 7A, 7B, “Starch”: “Bottom”). Starch accumulated in secondary phloem parenchyma in the treated AZ stems, while starch was not observed in the corresponding tissue of the control. The observed starch accumulation in the AZ supports the strong involvement of cytokinins in the development of the perennial stem with a storing function. Patterns of lipid bodies were similar between treated and untreated stems in cambium and phloem parenchyma (Figure 7A, 7B, “Lipids”).

**Figure 6:**
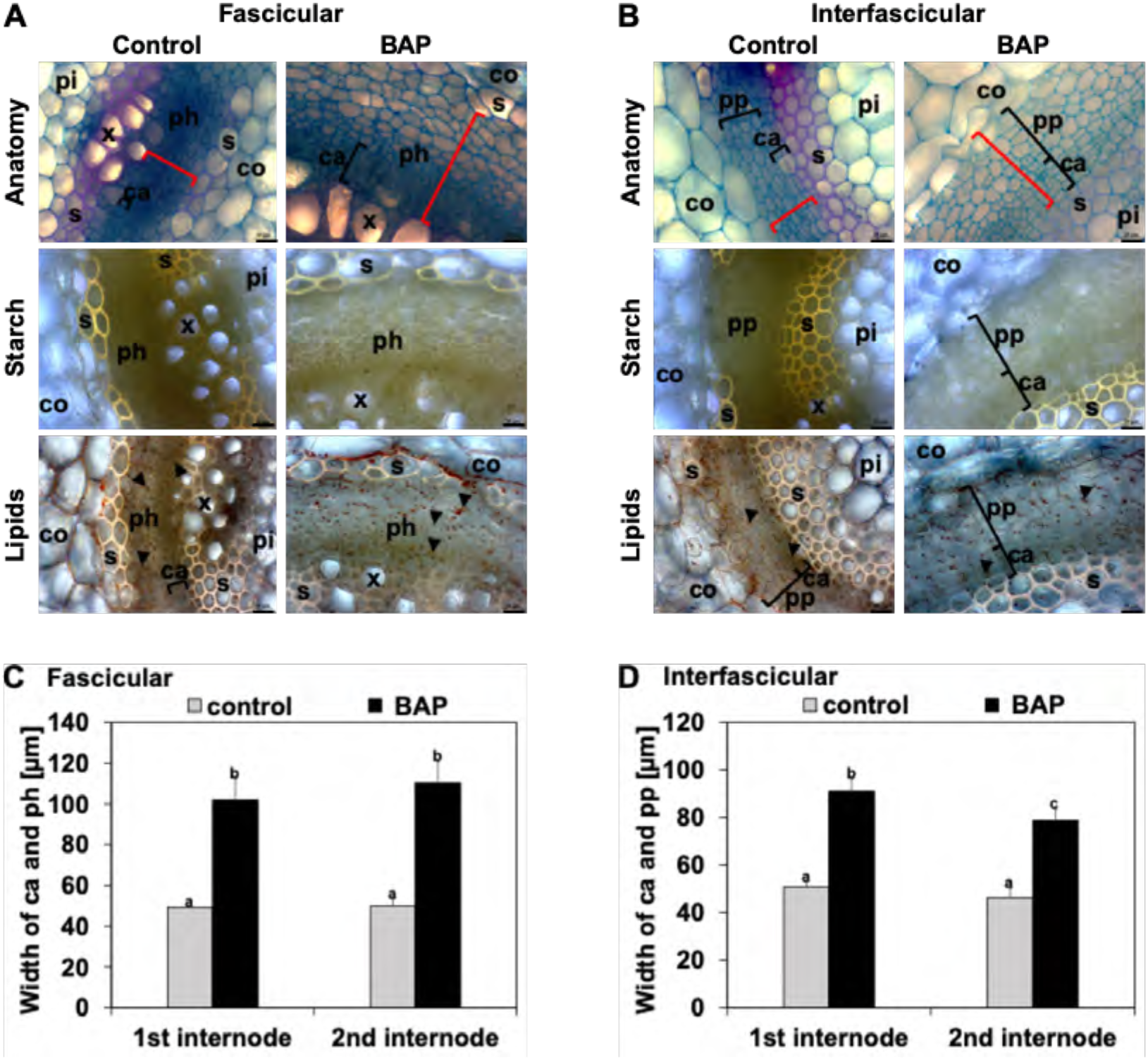
Effect of cytokinin treatment on cambium activity in the perennial zone (PZ). (A, B), Lateral stem first internode cross sections of (A), the fascicular region, (B), the interfascicular region of *A. alpina pep1-1*, following treatment with cytokinin (2 mM 6-benzylaminopurine, BAP) or mock control for eight days to the first internode. Representative images are shown. Anatomy, investigated following FCA staining, in blue, staining of non-lignified cell walls (parenchyma, phloem, meristematic cells), in red, staining of lignified cell walls and in greenish, staining of suberized cell walls (xylem, sclerenchyma, cork). Starch, visualized following Lugol’s iodine staining, dark violet-black color. Lipids, detected by Sudan IV staining, orange pinkish color of lipid bodies and yellowish color of suberized and lignified cell wall structures. Black triangles, lipid bodies; black brackets, responsive cambium and phloem/secondary phloem parenchyma; red brackets, quantified tissue width in (C) and (D). Abbreviations used in microscopic images: c, cork; ca, cambium; co, cortex; e, epidermis; pd, phelloderm; pe, periderm; ph, phloem including primary phloem, secondary phloem, phloem parenchyma; pi, pith; pp, secondary phloem parenchyma; s, sclerenchyma; x, xylem including primary xylem, secondary xylem, xylem parenchyma. Scale bars, 20 μm. (C, D) Quantified cambium and phloem/secondary phloem width in, (C), fascicular, and (D), interfascicular internode regions for first and second internodes. Data are represented as mean +/- SD (n = 3). Different letters indicate statistically significant differences, determined by one-way ANOVA-Tukey’s HSD test (p < 0.05).

**Figure 7:**
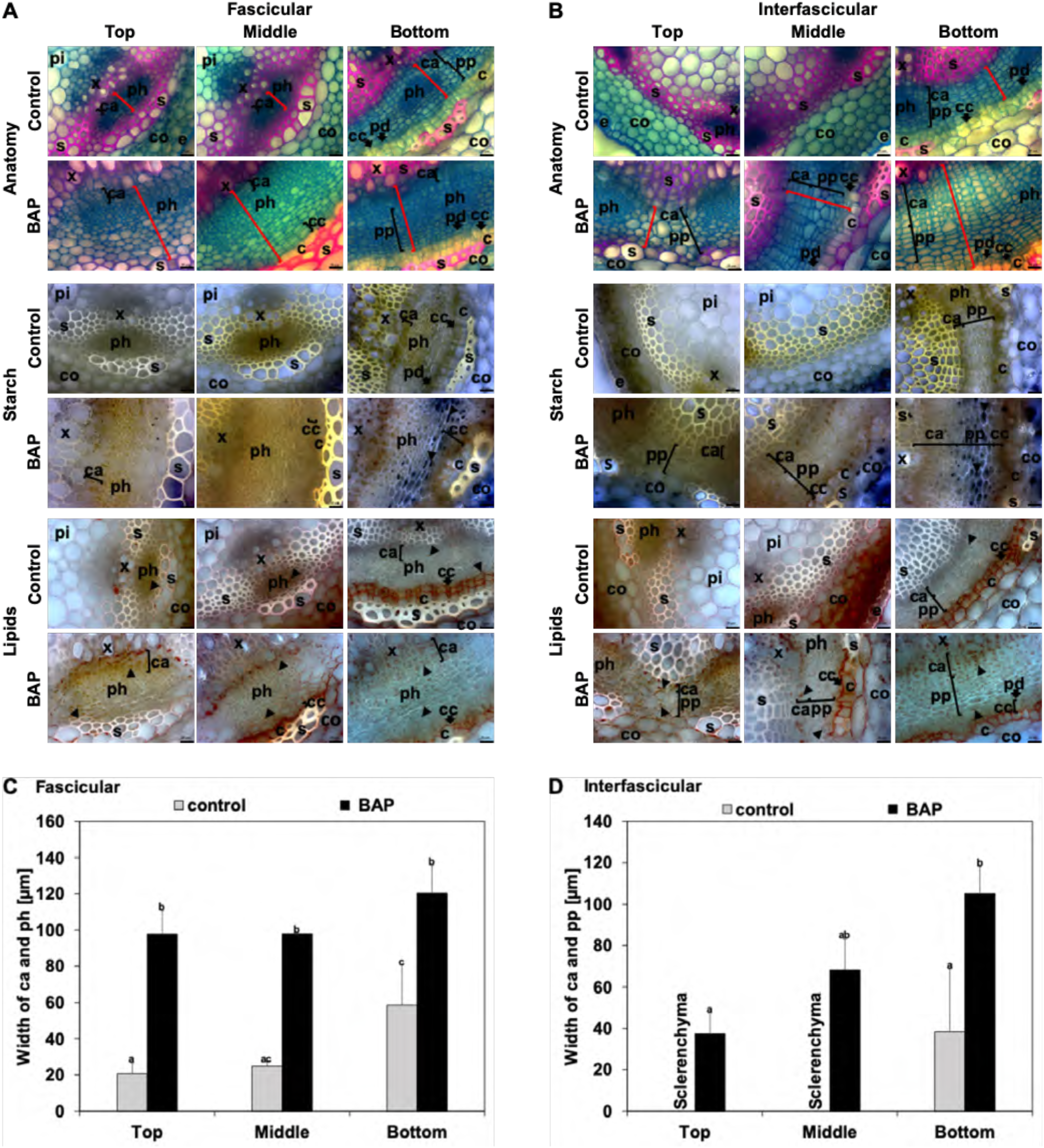
Effect of cytokinin treatment on cambium activity in the annual zone (AZ). (A, B), Lateral stem first internode cross sections of top, middle and bottom parts of the AZ (see also Supplemental Figure S5) in (A), the fascicular region, (B), the interfascicular region of *A. alpina perpetual flowering1-1* mutant (*pep1-1*), following treatment with cytokinin (2 mM 6-benzylaminopurine, BAP) or mock control for 46 days to all developing internodes of AZ. Representative images are shown. Anatomy, investigated following FCA staining, in blue, staining of non-lignified cell walls (parenchyma, phloem, meristematic cells), in red, staining of lignified cell walls and in greenish, staining of suberized cell walls (xylem, sclerenchyma, cork). Starch, visualized following Lugol’s iodine staining, dark violet-black color. Lipids, detected by Sudan IV staining, orange pinkish color of lipid bodies and yellowish color of suberized and lignified cell wall structures. Black triangles, starch and lipid bodies; black brackets, responsive cambium and phloem/secondary phloem parenchyma; red brackets, quantified tissue width in (C) and (D). Abbreviations used in microscopic images: c, cork; ca, cambium; co, cortex; e, epidermis; pd, phelloderm; pe, periderm; ph, phloem including primary phloem, secondary phloem, phloem parenchyma; pi, pith; pp, secondary phloem parenchyma; s, sclerenchyma; x, xylem including primary xylem, secondary xylem, xylem parenchyma. Scale bars, 20 μm. (C, D) Quantified cambium and phloem/secondary phloem width in, (C), fascicular, and (D), interfascicular internode regions in top, middle and bottom parts of the AZ. Data are represented as mean +/- SD (n = 3). Different letters indicate statistically significant differences, determined by one-way ANOVA-Tukey’s HSD test (p < 0.05).

Auxin (1-naphthaleneacetic acid, NAA), gibberellic acid (GA3) and ethylene precursor (1-aminocyclopropane-1-carboxylic acid, ACC) also influenced cambial activity in the first internode of the PZ upon application during eight days. Cambium and phloem parenchyma regions were increased in width, but in contrast to BAP to a maximum of 30 % and only in the treated first internode but not the second (Supplemental Figure S6, S7, S8). No changes with regard to starch and lipid body accumulation were observed (Supplemental Figure S6, S7, S8).

Taken together, among all tested hormones, cytokinin had the highest effect in the PZ on secondary growth. Cytokinin application to the AZ caused a shift in the zonation pattern and promoted secondary growth in the AZ, indicating that cytokinins act as signals to induce secondary growth. At an early growth stage, starch accumulation was not stimulated by cytokinins, whereas this was the case upon a prolonged treatment.

### Transcriptome analysis reflects cytokinin effects in the PZ

To provide support for PZ and AZ activities and obtain hints on signaling, we conducted a gene expression profiling experiment with *pep1-1* lateral stem internodes using RNA-seq at 4- and 5-week-old juvenile developmental PZ stages (stage I_PZ and stage II_PZ), a 7-week-old PZ stage with secondary growth and lipid bodies (stage III_PZ) and a 30-week-old mature PZ stage (stage IV_PZ) with advanced secondary growth and accumulated starch and lipid bodies and AZ internodes either in close proximity to the formed inflorescence (stage IV_AZ_if) or in the region with short internodes of the AZ (stage IV_AZ_si) (Figure 8A; Supplemental Table S2). Hierarchical clustering (HC) and Principal Component Analysis (PCA) confirmed the close relationships of three biological replicates of each sample and quality of RNA-seq data (Supplemental Figure S9). In HC, four distinct clusters were apparent, one cluster with stages IV_AZ_si and IV_AZ_if, a second one with stages I_PZ and II_PZ, grouping closely with AZ samples. The third and fourth cluster were stages III_PZ and IV_PZ, whereby the latter was most distant from all other groups (Supplemental Figure S9A). PCA analysis confirmed the expected variation between the samples. PC1 separated between the age of the internodes, similar as the first clusters in HC, while PC2 grouped according to PZ and AZ (Supplemental Figure S9B).

**Figure 8:**
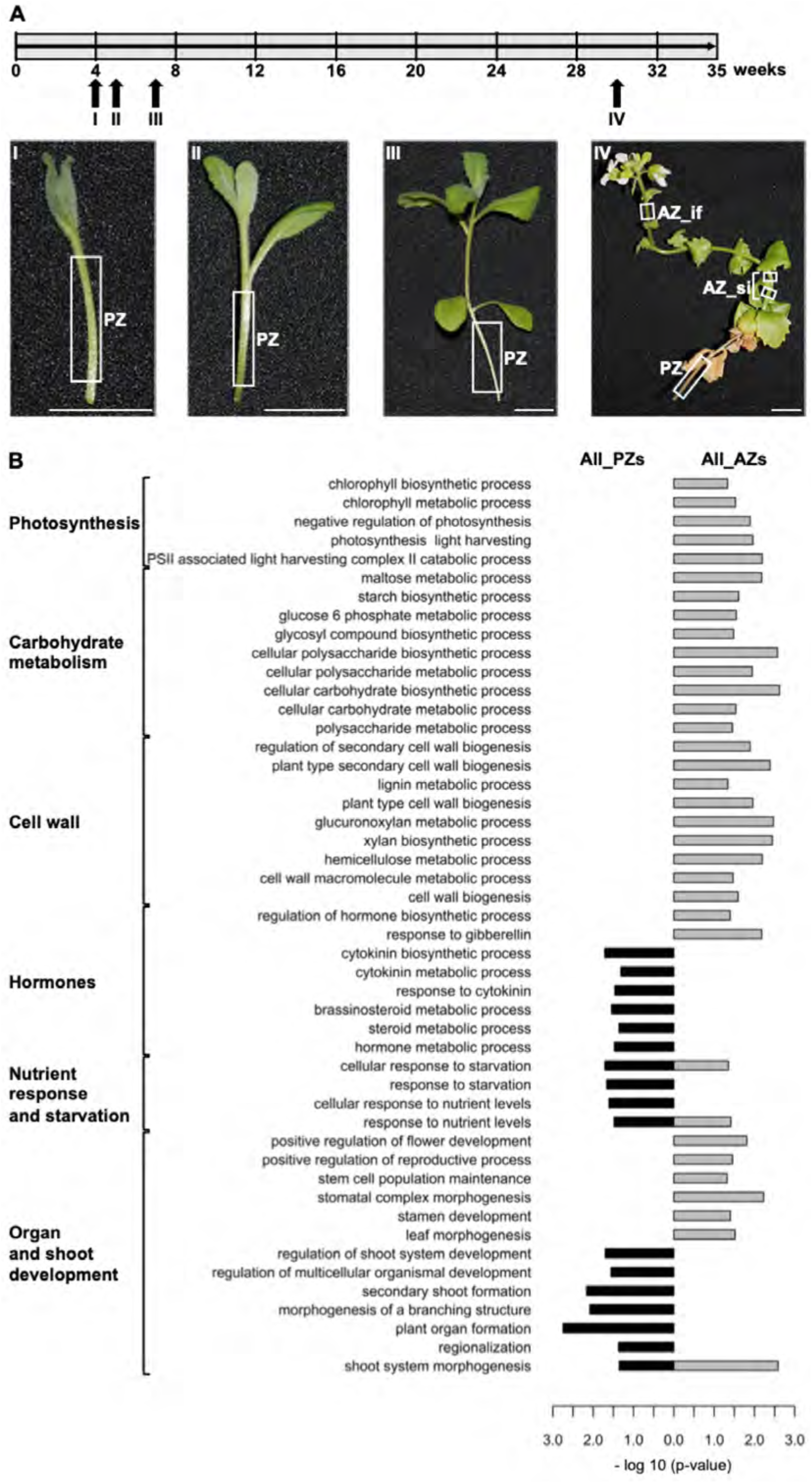
Transcriptome analysis of perennial (PZ) and annual (AZ) lateral stem zones. (A) Plant growth and morphology. Upper part, plant growth scheme and harvesting stages I-IV for RNA-sequencing (RNA-seq), long-day conditions at 20°C, *A. alpina perpetual flowering1-1* mutant (*pep1-1*). Bottom part, lateral stem morphology at each harvesting stage. Representative photos, white rectangles, harvested parts stage I_PZ and stage II_PZ, PZ both prior to visible secondary growth in a juvenile phase, stage III_PZ and stage IV_PZ, PZ with visible secondary growth, stage IV_AZ_si, small internode region, stage IV_AZ_if, inflorescence region. Scale bars, 1 cm. (B) GO term enrichment analysis of PZ versus AZ. PZ comprised the sum of stages I to IV_PZ, while AZ comprised stage IV_AZ_si and _if combined. GO terms were assigned to the indicated categories photosynthesis, carbohydrate metabolism, cell wall, hormones, nutrient response and starvation, organ and shoot development. Enriched, p < 0.05, represented as –log 10 values. Further information about the GO term enrichment analysis in Supplemental Table S18.

As we were seeking gene expression differences between PZ and AZ, we conducted meaningful crosswise enrichment analysis of gene ontology (GO) terms (Supplemental Tables S3-S17). Cytokinin terms were enriched in comparisons of stages II_PZ and IV_AZ_if, III_PZ and IV_AZ_si or IV_AZ_if (Supplemental Tables S11, S13, S14). By grouping the data and comparing all PZ with all AZ stages, enrichment analysis resulted in a total of 48 PZ- and 89 AZ-enriched GO terms, among them cytokinin-related terms (Figure 8B, Supplemental Table S18; confirmed by gene expression validation in Fig. 9). Other PZ-enriched GO terms were related to organ and shoot development, nutrient response and starvation genes, and other hormone responses (Figure 8B, Supplemental Table S18). AZ-enriched terms were cell wall, carbohydrate metabolism and photosynthesis categories (Figure 8B, Supplemental Table S18). GA biosynthesis and response terms were enriched in AZ (Figure 8B, Supplemental Table S18).

**Figure 9:**
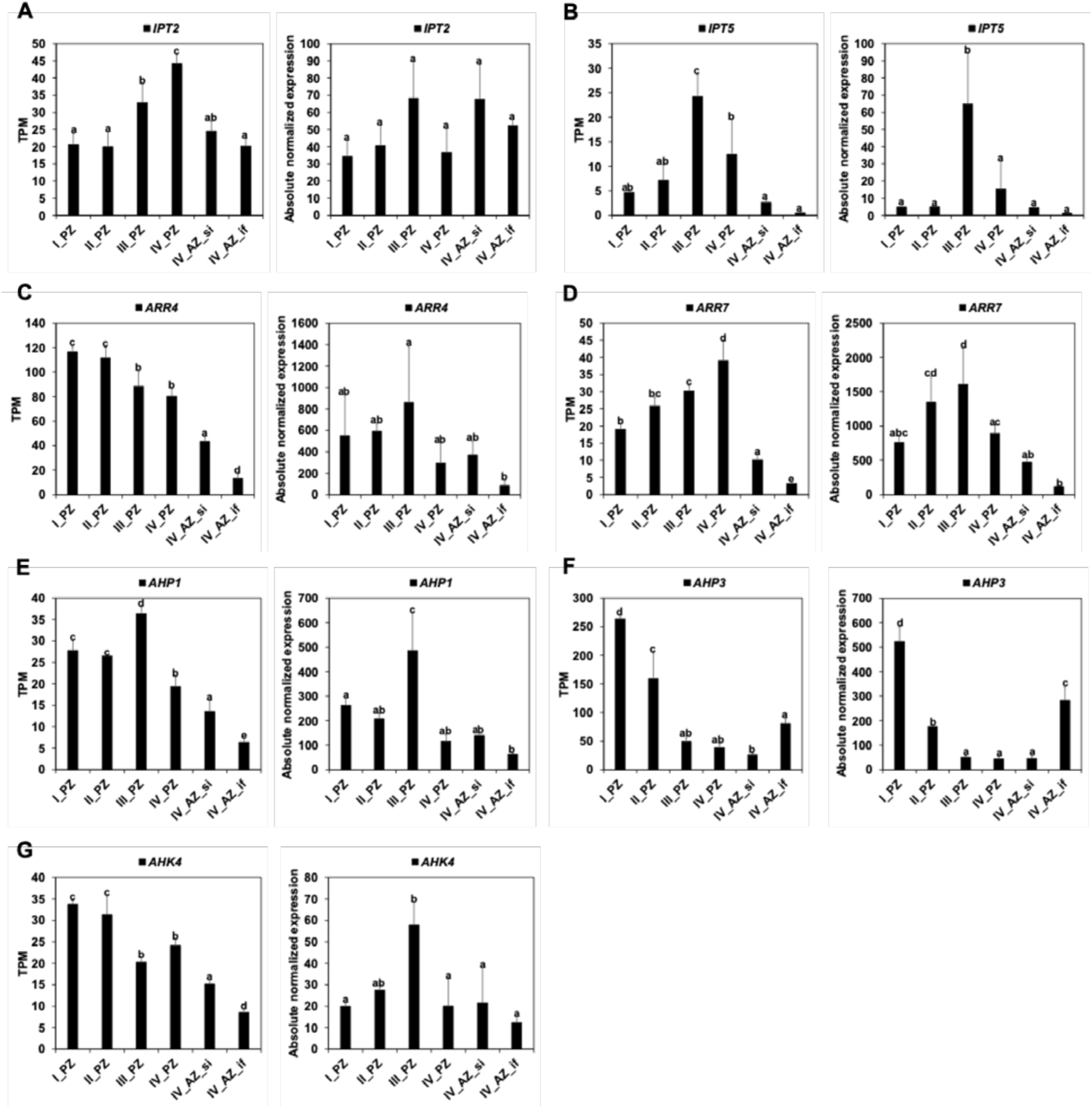
Gene expression of cytokinin signaling and response genes in lateral stem internodes. Gene expression was tested at different stages of the development of the perennial zone (PZ) and annual zone (AZ) in *A. alpina perpetual flowering1-1* mutant derivative (*pep1-1*. Represented are RNA-sequencing data (left; TPM, transcripts per million) and reverse transcription-qPCR data (right; Absolute normalized expression). (A), *IPT2, ISOPENTENYLTRANSFERASE 2*; (B) *IPT5, ISOPENTENYLTRANSFERASE 5*; (C) *ARR4, ARABIDOPSIS RESPONSE REGULATOR 4*; (D) *ARR7, ARABIDOPSIS RESPONSE REGULATOR 7*; (E) *AHP1, ARABIDOPSIS HISTIDINE-CONTAINING PHOSPHOTRANSMITTER 1*; (F) *AHP3, ARABIDOPSIS HISTIDINE-CONTAINING PHOSPHOTRANSMITTER 3*; (G) *AHK4, ARABIDOPSIS HISTIDINE KINASE 4*. Data are represented as mean +/- SD (n = 3). Different letters indicate statistically significant differences, determined by one-way ANOVA-Tukey’s HSD test (p < 0.05).

In summary, the PZ shows a high responsiveness to cytokinins as compared to the AZ, confirming that cytokinins play an important role in the differentiation of the PZ.

## Discussion

The complex perennial lifestyle of *A. alpina* comprises an allocation of nutrients and storage of high-energy C-compounds in the proximal vegetative perennial zone (PZ), characterized by secondary growth. The distal inflorescence annual zone (AZ) remains in a primary growth stage.

### *A. alpina* stems have proximal perennial (PZ) and distal annual zones (AZ) with C sequestration in the cambium and derivatives in the PZ

The vegetative growth zone (V) coincided with the PZ characterized by secondary growth, while the distal inflorescence zone (I) represented the AZ with primary growth. In the PZ-AZ transition zone secondary growth gradually shifted to primary growth. At the beginning of stem development, the anatomy of stem cross sections was similar in the PZ in *A. alpina* and AZ of *A. alpina* and *A. montbretiana*. With continuation of development, secondary growth was initiated by activities of cambium at first and then cork cambium. At the bottom of annual stems in *A. montbretiana* secondary growth was also found, where it may serve for general stability of upright stems, as described in *A. thaliana* (Agusti et al., 2011). Moreover, perennial life strategy is regarded to be ancestral (Hu et al., 2003; Grillo et al., 2009). Secondary growth at the bottom of *A. montbretiana* stems may still occur to some extent due to the inherited regulatory events. In the AZ, the interfascicular cambium and inner layer of the cortex differentiate into sclerenchyma during progression of development, perhaps for stabilization during flowering and silique production. The occurrence of PZ and AZ regions in all stems in *A. alpina* wild type Paj and *pep1-1* suggests that the genetically encoded signals for the PZ-AZ transition do not require vernalization treatment. Instead, we propose that a heterochronic regulation might control it. In the PZ, cambium and its derivatives store high-molecular weight C compounds in the form of starch and TAGs, in agreement with the rise in C contents and C/N ratios. The cell wall GO term was enriched in the AZ but not PZ, suggesting that the elevated C content in the PZ was not primarily due to cell wall biosynthesis. The enrichment of nutrient response and starvation GO terms in the PZ speaks in favor of a nutrient storage reservoir. Starch is present in the secondary phloem parenchyma of PZ and the outer cortex of AZ. This could be the reason why the GO term starch biosynthesis was not enriched in PZ. Starch deposition begins with the end of vegetative growth, usually coinciding with a cold treatment, however it is also in place in the absence of the cold treatment and increases up to the flowering stage and also during the senescence of leaves present in the PZ. C-containing compounds are presumably re-mobilized from senescing leaves to be allocated to adjacent storage tissue in the PZ. Interestingly, sequestration of C to phloem tissue of stems has been detected in trees, where functions are attributed to C supply during stress and in early spring (reviewed by Furze et al. 2018). In tree species storage also occurs in pith cells, wood ray and bark parenchyma (Netzer et al., 2018). It is thus not unusual that *A. alpina* uses different types of tissues for storage of different compounds. Lipid bodies occur in the cambium and cambium derivatives of the PZ and in cortex and phloem of the AZ, and this parallels generally a trend towards higher TAG content in the PZ than in the AZ. This suggests that lipids are used as storage compounds in the PZ. The N pool in *A. alpina* stems coincided with the protein pool, noteworthy in the early PZ but decreasing with development. Thus, N utilization in storage is not likely. Roots acted differently from stems and are not perennial starch storage organs in *A. alpina* but might be involved in lipid storage. In summary, C storage is more important than N storage, and the PZ of the stem has higher storage capacity than roots in *A. alpina*.

### Cytokinins are signals for demarcation of annual and perennial growth zones

The open question is which events are responsible to demarcate the sharp PZ-AZ transition. The PZ-AZ boundary separates besides secondary/primary growth and C storage also organ differentiation at the nodes and in axils, such as bud outgrowth into fully developed lateral stems, bud dormancy, bud differentiation into a singular flower or bud outgrowth into flowering secondary and tertiary branches. The small PZ-AZ transition zone has short internodes and is marked by a gradual shift from secondary to primary growth. Multiple signals could act in these differentiation processes in the PZ and AZ.

One possible explanation is that the PZ state is an initial ground state. In this scenario, the progression to the PZ becomes halted later in developmental time in the AZ. In this model, the PZ to AZ transition is a heterochrony or developmental phase transition. Since regulation of bud fate is under control of meristem identity genes, mutants in those genes affect developmental phase transitions. In annuals, some progression to the PZ may take place in a rudimentary manner at the bottom of stems or during senescence onset for stabilization reasons, however, a full progression to the PZ is prevented. We exclude that flowering or vernalization emit signals for the PZ-AZ transition in Paj and *pep1-1*. The *pep1-1* mutant has also the PZ-AZ transition. Even though the presence of flowers was not a requirement for PZ-AZ transition, upstream regulators of flowering and developmental phase transition, namely the regulatory network of microRNA miR156, SPL proteins, and Mediator complex subunits may well control aspects of stem differentiation (Park et al. 2017; Bergonzi et al. 2013; Guo et al. 2017; Hyun et al. 2019; Zhang and Guo, 2020).

Developmental decisions are often taken under the influence of plant hormones and they affect stem differentiation, flowering and developmental phase transition. Cytokinin application shifted the PZ-AZ transition and might be coupled to a heterochronic signal. A cytokinin effect was further confirmed by enrichment of cytokinin biosynthesis and response gene expression in the developing PZ. Even application directly to AZ caused an extension of PZ into AZ. Thus, cytokinins might be signals that promote the progression during developmental phase transitions, and cytokinin depletion rather than decreased sensitivity might trigger the precocious halting in AZ. Cytokinins play also a crucial role in the regulation of source and sink relationships including starch deposition (Thomas, 2013). Cytokinins are likely signals in the PZ to promote secondary growth and later in combination with flowering and senescence also sink activity and starch formation. Cytokinins also retard senescence which is partly explained by cytokinin-modified source-sink relationships (Thomas, 2013). Together, this suggests that cytokinins are less likely to act in the AZ but rather contribute to the maintenance of an adult vegetative state in the PZ.

GA, auxin and ethylene also promoted secondary growth in PZ. Yet, these hormones acted only locally on the same internode and not to the same extent as cytokinin. Auxin produced in V3 lateral branches may cause bud dormancy in V2, according to an auxin canalization model. This growth inhibition effect is also under the control of vernalization and further promoted by enhanced sinks for nutrients in the AZ and V3 lateral branches (Vayssières et al. 2020). In our experiments, auxin did not stimulate secondary growth in the neighboring internodes, that had not been treated, in the way that cytokinin did. Auxin signaling may give rise to distinctive V subzones in the PZ but not the PZ-AZ transition. Gibberellins (GAs) are influential in the xylem region promoting xylem cell differentiation and lignification (Denis et al., 2017). Interestingly, this stronger lignification of vascular cells resembles the anatomy which has been detected in our study in the AZ of Paj and *pep1-1* as well as in *A. montbretiana* at later developmental stages. Perhaps GA signals explain the sensitivity of cambium in AZ to develop lignified cell walls and differentiate into sclerenchyma. Indeed, GA-related GO terms were enriched in AZ versus PZ, supporting this idea.

To test whether hormone gradients and developmental phase transition regulation are responsible for the secondary to primary growth transition, transgenic approaches will be helpful in the future to manipulate these responses in the different growth zones.

## Conclusions

The functional complexity of stem differentiation in *Arabis* offers the possibility to study regulation of perennial (advanced secondary growth and storage) and annual (primary growth and senescence) traits using the same model species. The close relationship among *Brassicaceae* and the similarities of secondary growth processes in trees and *Arabidopsis* can be exploited to unveil signaling pathways and regulators for stem differentiation in *A. alpina* in future projects. Further studies need to show whether heterochronic PZ-AZ transition genes and plant hormones determine the transport of effectors, the sensitivity to respond to effectors or to alter specific long-distance signaling processes, and how these processes link flowering control and secondary growth.

## Acknowledgements

We acknowledge the excellent technical assistance of M. Graf, E. Klemp and K. Weber for GC-MS and EA-IRMS measurements. We thank Vera Wewer (University of Cologne) for her advice regarding the establishment of the lipid extraction protocol. The authors thank E. Wieneke for help with gene expression studies. We are grateful to Maria C. Albani, University of Cologne, for critical reading of the manuscript. This project was funded by the Deutsche Forschungsgemeinschaft (DFG) under Germany’s Excellence Strategy – EXC 2048/1 – project 390686111. H.L. was the recipient of a doctoral fellowship from Northwest A&F University, College of Horticulture, Yangling, China.

