## Supplemental figures for "Cytokinin-promoted secondary growth and nutrient storage in the perennial stem zone of *Arabis alpina*"

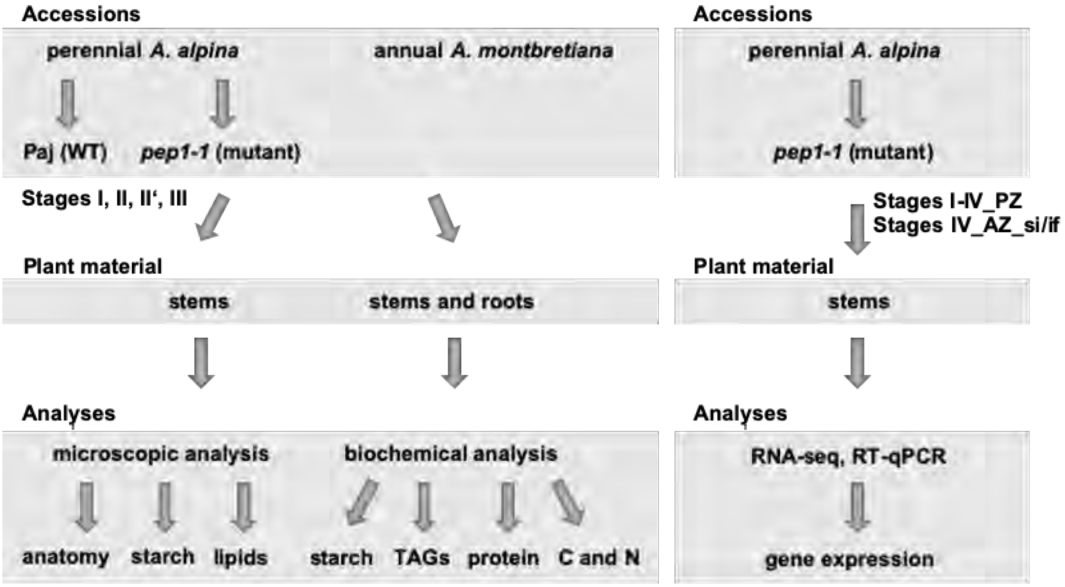


**Supplemental Figure S1:** Experimental outline. Abbreviations: perennial *A. alpina* Pajares (Paj WT, wild type); its *perpetual flowering1-1* mutant derivative (*pep1-1*); perennial zone (PZ); annual zone (AZ); short internode zone (si); inflorescence zone (if); triacylglycerol (TAG); carbon (C); nitrogen (N); RNA-sequencing (RNA-seq); reverse transcription-qPCR (RT-qPCR).

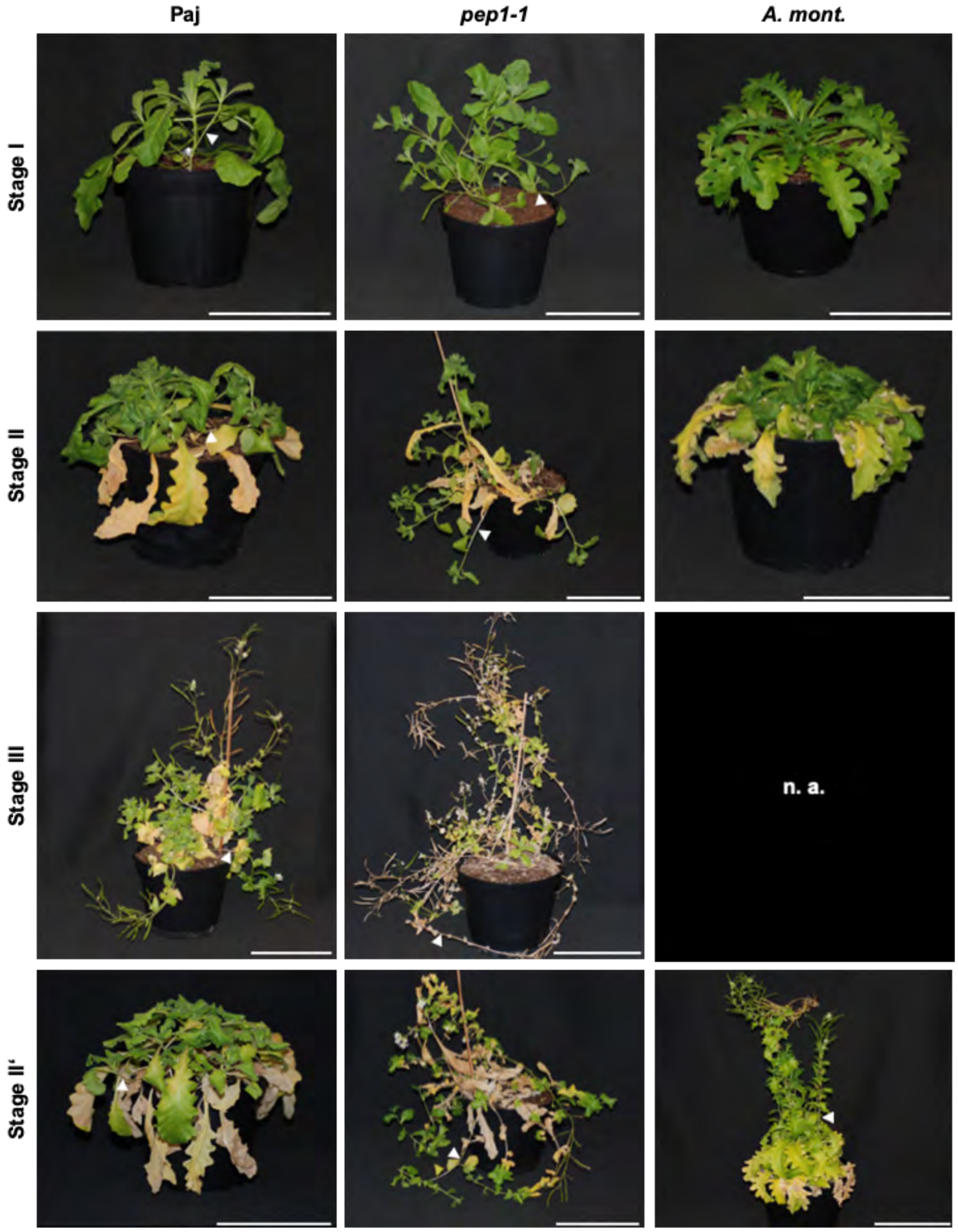


**Supplemental Figure S2:** Whole plant morphology. Representative photos of plants at stage I, stage II, stage III, stage II' for microscopic and biochemical analysis; perennial *A. alpina* Pajares (Paj, wild type), its *perpetual flowering1-1* mutant derivative (*pep1-1*) and annual *A. montbretiana* (*A. mont.*). White triangles indicate representative lateral stems used analyses (see Figure 2); n. a., plants not available at this stage. Scale bar, 10 cm.

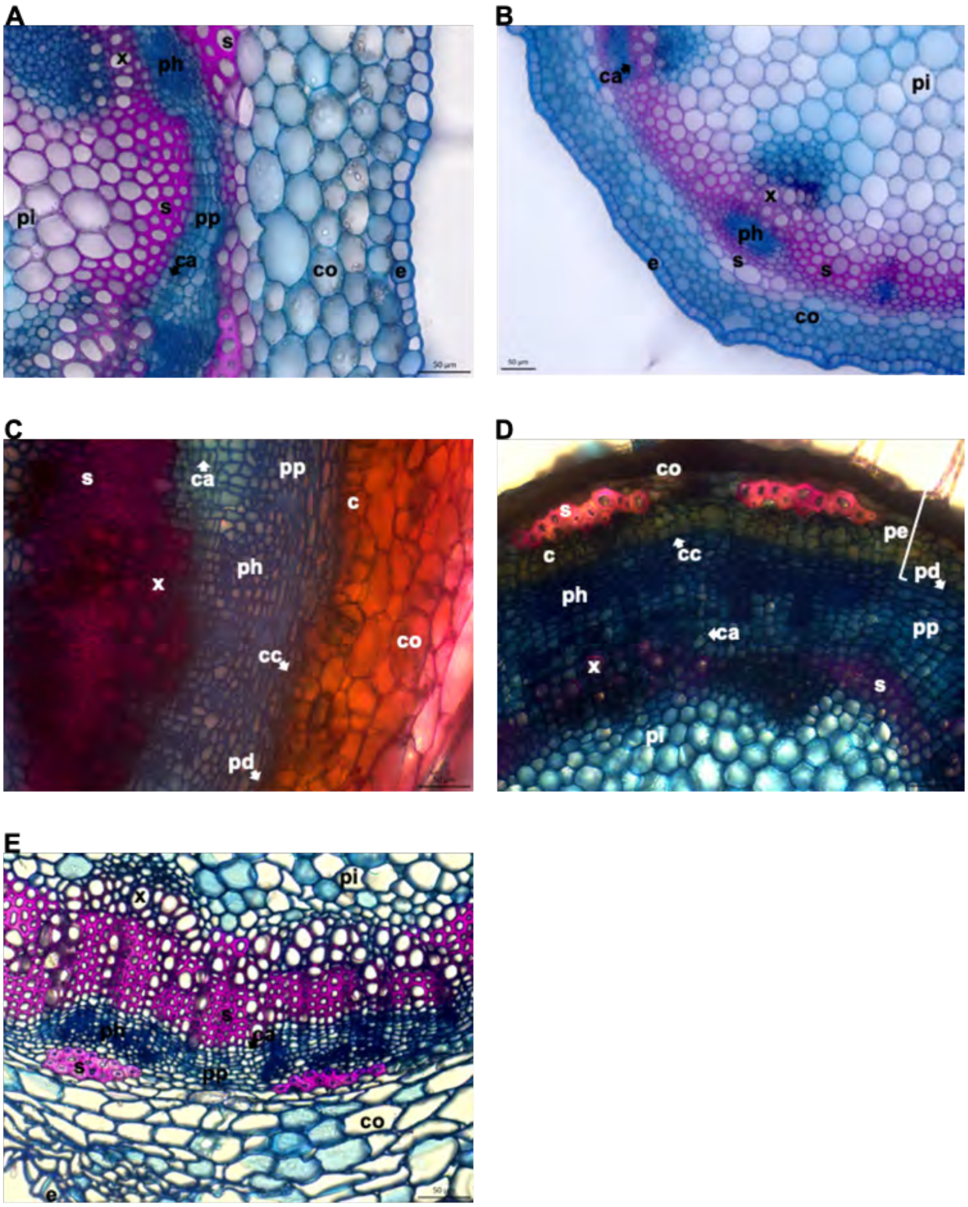


**Supplemental Figure S3:** Additional internode cross section analysis. (A) Early stage of development in between stages II and III of the upper annual stem zone of *perpetual flowering1-1* mutant; the center of the image shows the interfascicular region. (B) Late stage of development beyond stage III of the annual stem zone of wild type Pajares (Paj). (C, D), Transition regions at stage III in (C), Paj, and (D), *pep1-1*. (E) Internode region of *A. montbretiana* at stage II' close to the main axis. Abbreviations used in microscopic images: c, cork; ca, cambium; co, cortex; e, epidermis; pd, phelloderm; pe, periderm; ph, phloem including primary phloem, secondary phloem, phloem parenchyma; pi, pith; pp, secondary phloem parenchyma; s, sclerenchyma; x, xylem including primary xylem, secondary xylem, xylem parenchyma. Arrows point to respective tissues. Scale bars, 50 µm.

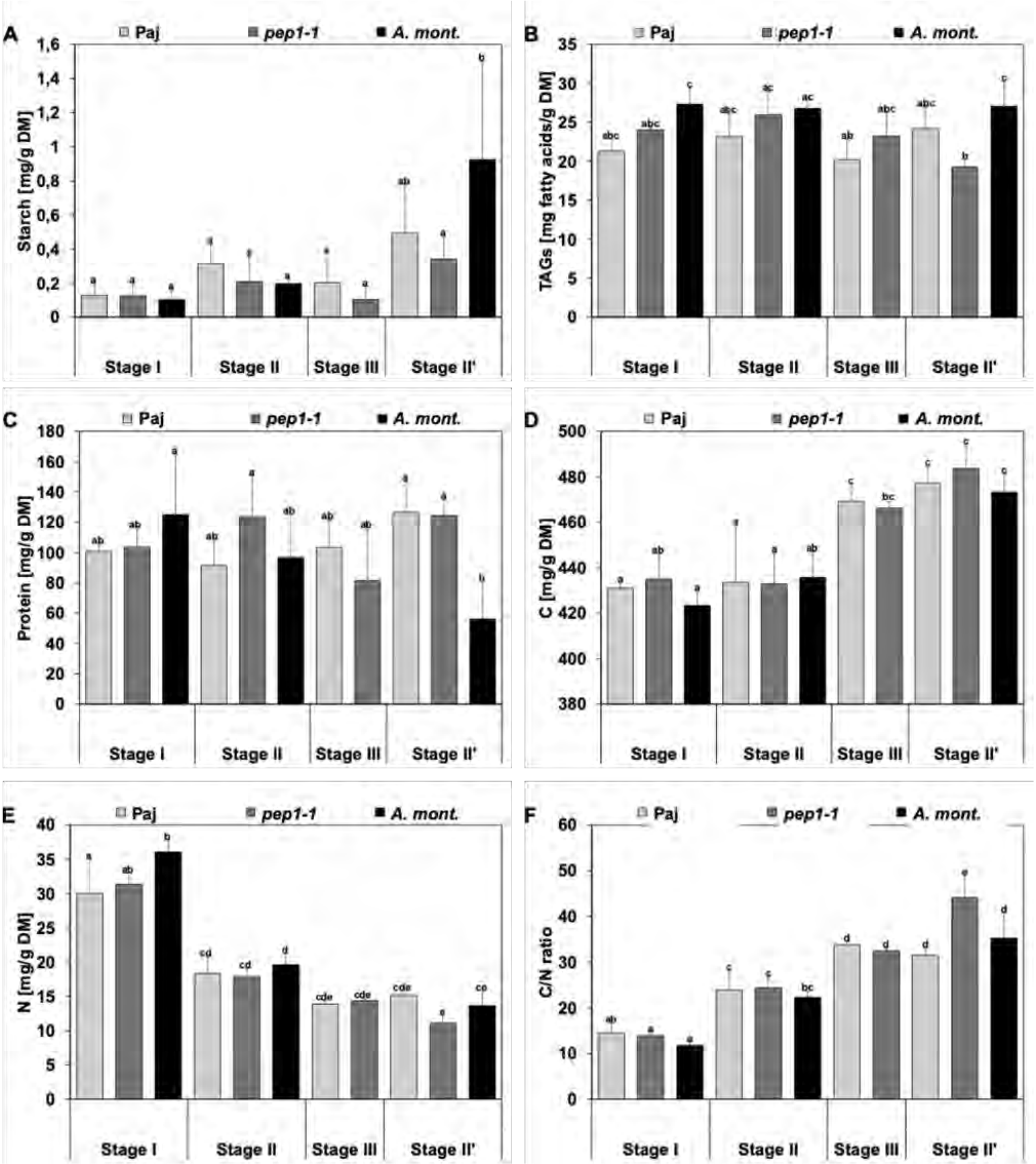


**Supplemental Figure S4:** Analysis of nutrient storage in roots. Stem diagrams represent contents per dry matter (DM) of (A), starch, (B), triacylglycerols (TAGs), (C), protein, (D), carbon (C), (E), nitrogen (N), (F), C/N ratios. Entire root systems were harvested at stages I, II, III and II' of *A. alpina* Pajares (Paj, wild type), its *perpetual flowering1-1* mutant derivative (*pep1-1*) and annual *A. montbretiana* (*A. mont.*). Data are represented as mean +/- SD (n = 3-7). Different letters indicate statistically significant differences, determined by one-way ANOVA-Tukey's HSD test ($p < 0.05$).

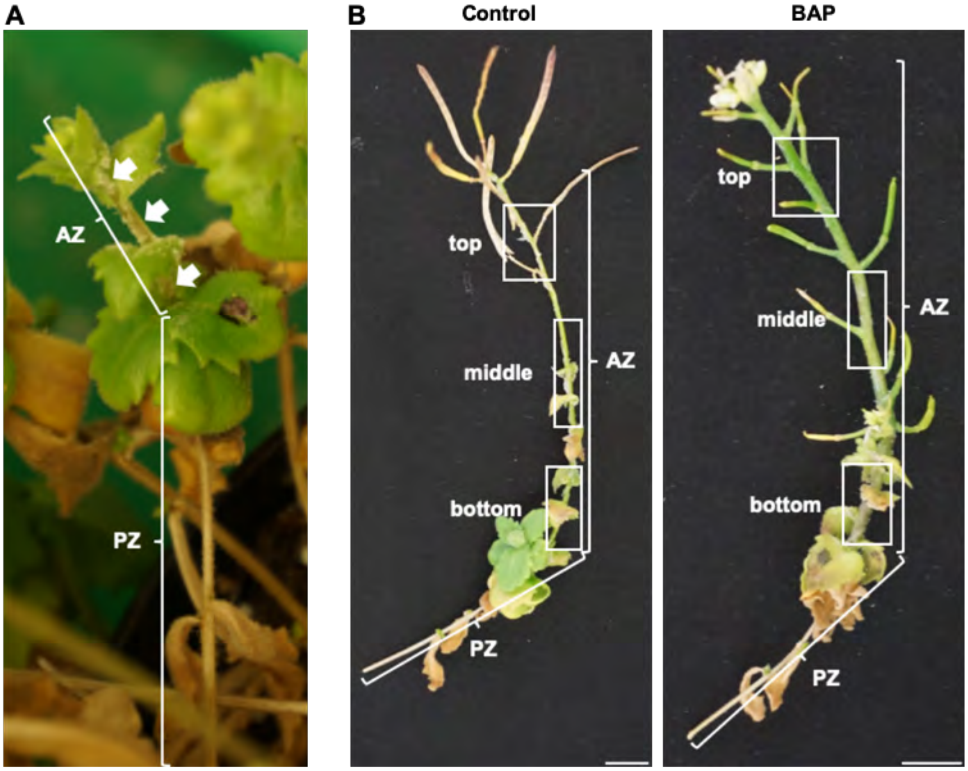


**Supplemental Figure S5:** Effect of prolonged cytokinin treatment and dissection of the annual zone (AZ). (A) Representative lateral branch of *perpetual flowering1-1* mutant before treatment. Three internodes belonging to the AZ were treated on day 1 (arrows) with 2 mM 6-benzylaminopurine (BAP) or mock treatment. These and following newly formed internodes were treated every fourth day until the experiment was stopped after 46 days. (B) Representative lateral branches after 46-day BAP and mock treatment (control). For the microscopic analysis, the AZ was divided into three regions (boxes: top, middle and bottom), which were comparable between control and cytokinin treatment with regard to the overall size of the formed AZ. Each investigated region comprised two to three internodes.

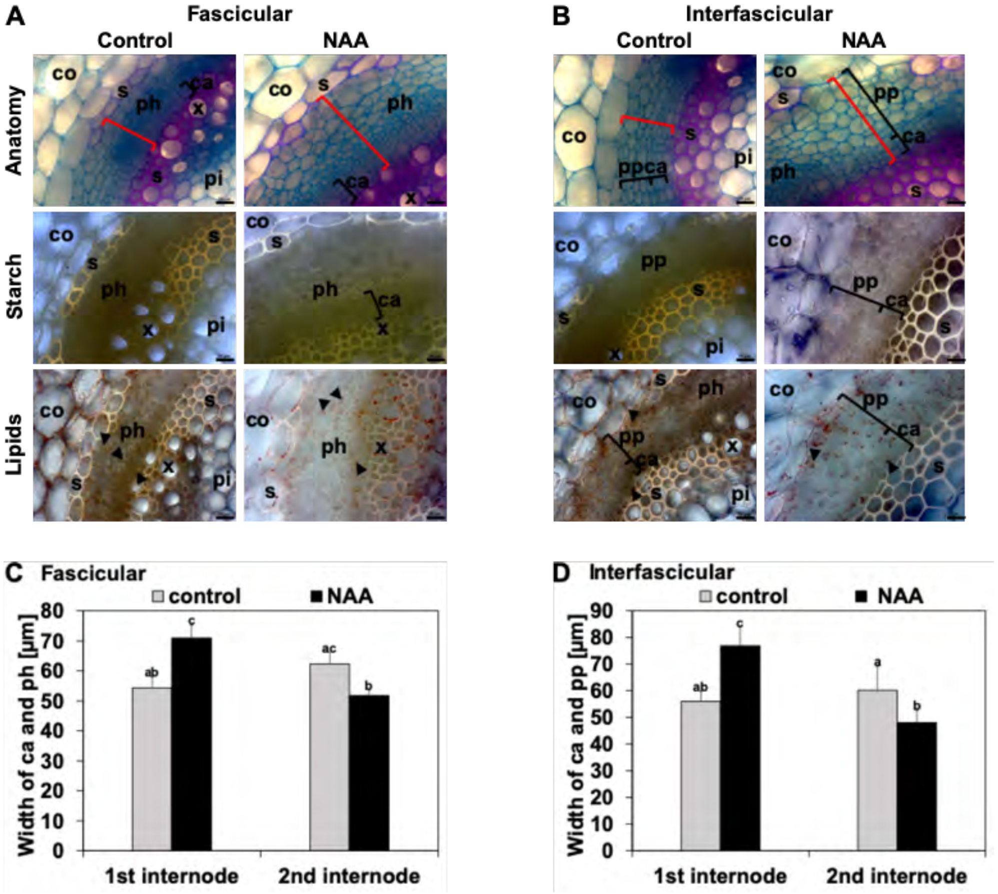


**Supplemental Figure S6:** Effect of auxin treatment on cambium activity in the perennial zone (PZ). (A, B), Lateral stem first internode cross sections of (A), the fascicular region, (B), the interfascicular region of *A. alpina perpetual flowering1-1* mutant, following treatment with auxin (0.5 mM 1-naphthaleneacetic acid, NAA) or mock control for eight days to the first internode. Representative images are shown. Anatomy, investigated following FCA staining, in blue, staining of non-lignified cell walls (parenchyma, phloem, meristematic cells), in red, staining of lignified cell walls and in greenish, staining of suberized cell walls (xylem, sclerenchyma, cork). Starch, visualized following Lugol's iodine staining, dark violet-black color. Lipids, detected by Sudan IV staining, orange pinkish color of lipid bodies and yellowish color of suberized and lignified cell wall structures. Black triangles, lipid bodies; black brackets, responsive cambium and phloem/secondary phloem parenchyma; red brackets, quantified tissue width in (C) and (D). Abbreviations used in microscopic images: c, cork; ca, cambium; co, cortex; e, epidermis; pd, phelloderm; pe, periderm; ph, phloem including primary phloem, secondary phloem, phloem parenchyma; pi, pith; pp, secondary phloem parenchyma; s, sclerenchyma; x, xylem including primary xylem, secondary xylem, xylem parenchyma. Scale bars, 20 µm. (C, D) Quantified cambium and phloem/secondary phloem width in, (C), fascicular, and (D), interfascicular internode regions for first and second internodes. Data are represented as mean +/- SD (n = 5). Different letters indicate statistically significant differences, determined by one-way ANOVA-Tukey's HSD test ($p < 0.05$).

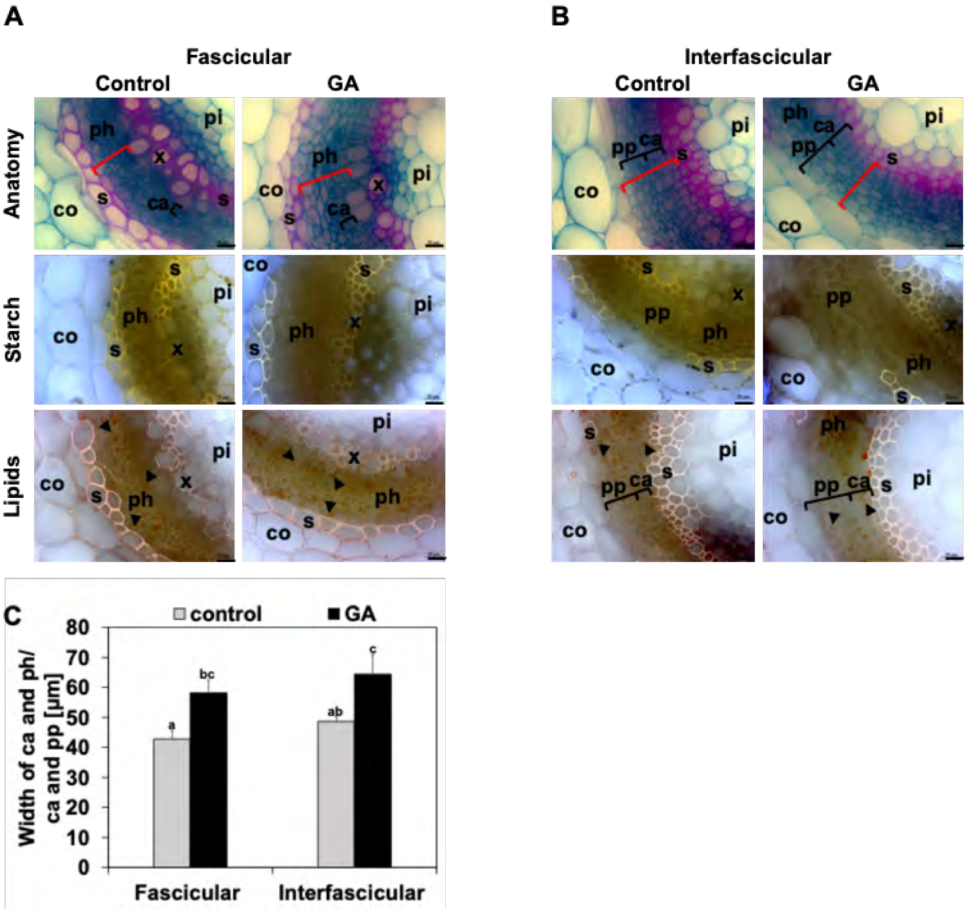


**Supplemental Figure S7:** Effect of gibberellic acid (GA) treatment on cambium activity in the perennial zone (PZ). (A, B), Lateral stem first internode cross sections of (A), the fascicular region, (B), the interfascicular region of *A. alpina perpetual flowering1-1* mutant, following treatment with GA (4 mM gibberellin $A_3$) or mock control for eight days to the first internode. Representative images are shown. Anatomy, investigated following FCA staining, in blue, staining of non-lignified cell walls (parenchyma, phloem, meristematic cells), in red, staining of lignified cell walls and in greenish, staining of suberized cell walls (xylem, sclerenchyma, cork). Starch, visualized following Lugol's iodine staining, dark violet-black color. Lipids, detected by Sudan IV staining, orange pinkish color of lipid bodies and yellowish color of suberized and lignified cell wall structures. Black triangles, lipid bodies; black brackets, responsive cambium and phloem/secondary phloem parenchyma; red brackets, quantified tissue width in (C). Abbreviations used in microscopic images: c, cork; ca, cambium; co, cortex; e, epidermis; pd, phelloderm; pe, periderm; ph, phloem including primary phloem, secondary phloem, phloem parenchyma; pi, pith; pp, secondary phloem parenchyma; s, sclerenchyma; x, xylem including primary xylem, secondary xylem, xylem parenchyma. Scale bars, 20 µm. (C) Quantified cambium and phloem/secondary phloem width in fascicular and interfascicular internode regions for first internode. Data are represented as mean +/- SD (n = 3). Different letters indicate statistically significant differences, determined by one-way ANOVA-Tukey's HSD test ($p < 0.05$).

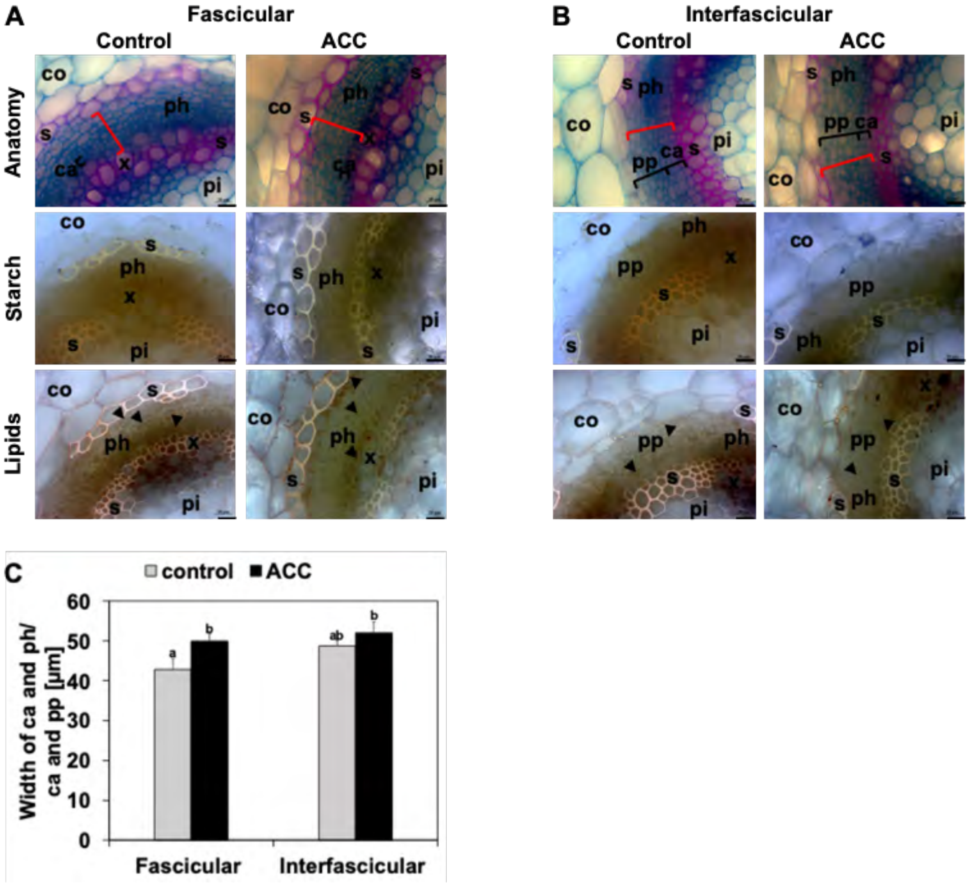


**Supplemental Figure S8:** Effect of ethylene on cambium activity in the perennial zone (PZ). (A, B), Lateral stem first internode cross sections of (A), the fascicular region, (B), the interfascicular region of *A. alpina perpetual flowering1-1* mutant, following treatment with 2 mM 1-aminocyclopropane-1-carboxylic acid (ACC) or mock control for eight days to the first internode. Representative images are shown. Anatomy, investigated following FCA staining, in blue, staining of non-lignified cell walls (parenchyma, phloem, meristematic cells), in red, staining of lignified cell walls and in greenish, staining of suberized cell walls (xylem, sclerenchyma, cork). Starch, visualized following Lugol's iodine staining, dark violet-black color. Lipids, detected by Sudan IV staining, orange pinkish color of lipid bodies and yellowish color of suberized and lignified cell wall structures. Black triangles, lipid bodies; black brackets, responsive cambium and phloem/secondary phloem parenchyma; red brackets, quantified tissue width in (C). Abbreviations used in microscopic images: c, cork; ca, cambium; co, cortex; e, epidermis; pd, phelloderm; pe, periderm; ph, phloem including primary phloem, secondary phloem, phloem parenchyma; pi, pith; pp, secondary phloem parenchyma; s, sclerenchyma; x, xylem including primary xylem, secondary xylem, xylem parenchyma. Scale bars, 20 µm. (C) Quantified cambium and phloem/secondary phloem width in fascicular and interfascicular internode regions for first internode. Data are represented as mean +/- SD (n = 3). Different letters indicate statistically significant differences, determined by one-way ANOVA-Tukey's HSD test ($p < 0.05$).

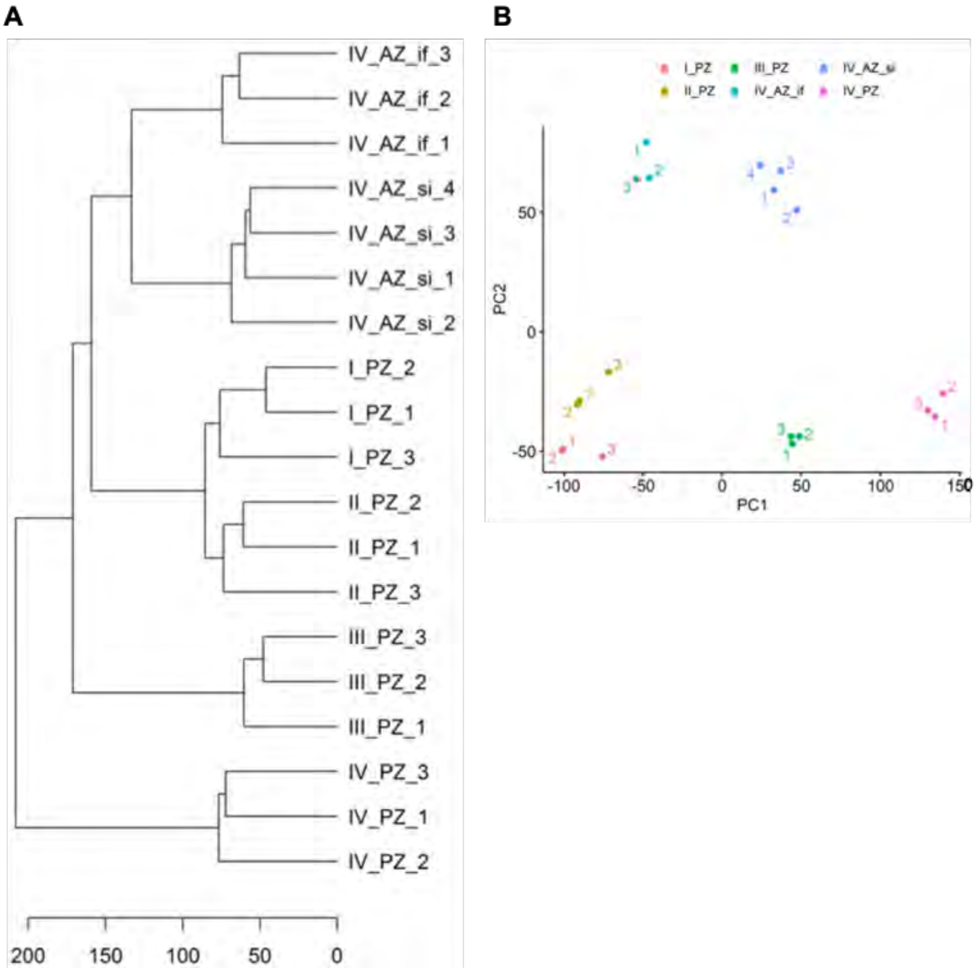


**Supplemental Figure S9:** Hierarchical clustering (HC) and Principal Component Analysis (PCA) of RNA-sequencing data. (A) HC and (B) PCA corresponding to internode samples harvested at different stages from perennial (PZ) and annual (AZ) lateral stem zones, stage I_PZ, stage II_PZ, stage III_PZ, stage IV_PZ, stage IV_AZ_si, stage IV_AZ_if; biological replicates 1-3 or 1-4 (compare with Figure 8A).
